# Single-cell profiling reveals a novel RAB13+ endothelial subpopulation and profibrotic mesenchymal cells in the aged human bone marrow

**DOI:** 10.1101/2025.01.28.635238

**Authors:** Itziar Cenzano, Miguel Cócera, Ana Rosa Lopez-Perez, Lorea Campos-Dopazo, Javier Ruiz, Ignacio Sancho, Patxi San Martin-Uriz, Paula Aguirre-Ruiz, Sarai Sarvide, Amaia Vilas-Zornoza, Purificacion Ripalda-Cemborain, Diego Alignani, Aitziber Lopez, Marta Miñana Barrios, Delia Quilez Agreda, Jin Ye, Robert Lehmann, Laura Sudupe, Marta Abengozar-Muela, Luis-Esteban Tamariz-Amador, Emma Muiños-López, Borja Saez, Jesper Tegner, Isabel A. Calvo, David Gomez-Cabrero, Felipe Prosper

**Author notes:** These authors contributed equally to this work (Itziar Cenzano, Miguel Cócera). These authors jointly supervised this work (Itziar Cenzano, Isabel A. Calvo, David Gomez-Cabrero, Felipe Prosper).

## Abstract

The bone marrow (BM) microenvironment plays a crucial role in regulating hematopoiesis, yet the molecular changes associated with aging in humans remain poorly understood. Using single-cell RNA sequencing, we uncovered transcriptional shifts in BM endothelial cells (EC) and mesenchymal stem cells (MSC) during aging. Aged sinusoidal EC exhibited a prothrombotic phenotype with compromised mitochondrial and vascular function. Additionally, we identified a novel arterial EC subset, emerging in aged individuals, characterized by RAB13 expression and associated with transcriptional regulatory processes. MSC from aged subjects displayed impaired matrix remodeling and epithelial-mesenchymal transition, driven partly by a subpopulation of THY1+ profibrotic cells absent in younger individuals. Finally, immunofluorescent imaging and spatial transcriptomics confirmed the presence of these aging-associated cells in BM samples from aged individuals. In summary, this work provides a comprehensive view of the transcriptional landscape, cellular interactions, and spatial organization of aged EC and MSC, offering novel insights and potential targets that could be exploited for preventing age-associated changes in humans.

**Teaser:** Aging reshapes the bone marrow with emergence of RAB13+ endothelial cells and profibrotic stromal cells altering tissue function.

## Introduction

Hematopoietic stem cells (HSC) are the fundamental pillar of the hematopoietic system, sustaining blood production for the entire lifespan of an organism(*1*). The HSC reside within a specialized bone marrow (BM) microenvironment called niche(*2*). The HSC niche consists of a complex and dynamic molecular network of interactions among hematopoietic and non-hematopoietic cells, extracellular matrix (ECM) components, and soluble factors, all working together to ensure proper hematopoiesis(*3–5*).

Aging induces a progressive decline of the anatomical and physiological function of all organ systems(*6*) and is the major risk factor across different diseases. In the hematopoietic system, aging is characterized by an increased number of HSC accompanied by reduced self-renewal capacity and regenerative potential(*7*). Over the past decades, several molecular mechanisms that control the HSC aging process have been revealed, yet our understanding remains incomplete(*8*). What has been established is that the senescent phenotype results from the combination and interplay of the intrinsic aging of the HSC and the detrimental effects of an aged microenvironment(*8*, *9*). The study of the microenvironment has demonstrated that aging causes a reduction in the number of endothelial cells (EC), significant disorganization of vascular structures, decreased angiogenic potential, and increased vascular leakiness and reactive oxygen species (ROS)(*10–12*). Additionally, aged mesenchymal stromal cells (MSC) exhibit impaired differentiation into osteoblasts towards adipocytes and secretion of pro-inflammatory cytokines influencing the skewed myeloid differentiation of aged HSC(*13*, *14*). Therefore, to fully comprehend and possibly mitigate the effects of aging in the hematopoietic system, it is necessary to dissect the complex interactions between aged HSC and their aged non-hematopoietic microenvironment.

In that direction, recent studies have shed light on remodeling the non-hematopoietic BM microenvironment in mice during aging(*15*, *16*). However, translating these findings to humans has been challenging, primarily due to difficulties in adapting methodologies from mouse models and obtaining high-quality human samples with adequate EC and MSC populations(*17–21*). Consequently, these limitations have restricted the scope of single-cell analyses in the non-hematopoietic BM compartment.

In this article, we profiled EC and MSC at both single-cell and spatial resolution from human BM samples of young and elderly healthy donors. Our analysis reveals that aged sinusoidal EC display a prothrombotic profile and impaired mitochondrial and vascular functions. We identify an aged arterial-like EC subset marked by RAB13 expression, which is linked to dysregulated transcriptional elongation activity. In the stromal compartment, aging drives matrix remodeling mediated partly by THY1+ profibrotic stromal cells, increases oxidative phosphorylation (OXPHOS), and shows a reduced unfolded protein response. Furthermore, we explore the spatial context of these alterations using immunofluorescent imaging and spatial transcriptomics. These spatial approaches confirm the presence of RAB13+ EC in aged samples and uncover significant age-related disruptions in cell-cell communication within the BM niche. This comprehensive analysis offers new insights into the transcriptomic alterations that occur during aging in the human BM.

## Results

### Transcriptional profiling of human young and aged BM endothelial and stromal cells

To investigate age-related changes in the human non-hematopoietic BM microenvironment, we performed single-cell transcriptomics on prospectively fluorescence-activated cell-sorted EC and MSC from BM tissue of healthy young (<50 years of age; mean 37,8) and aged (>58 years of age; mean 68,5) individuals undergoing orthopedic surgery (**Fig. 1a, Fig. S1 and Table S1**). After initial quality control and pre-processing, we obtained a total of 35368 cells across all samples (14283 from young and 21085 from aged individuals), which, despite our sorting strategy, included contaminant hematopoietic cells.

**Figure 1.**
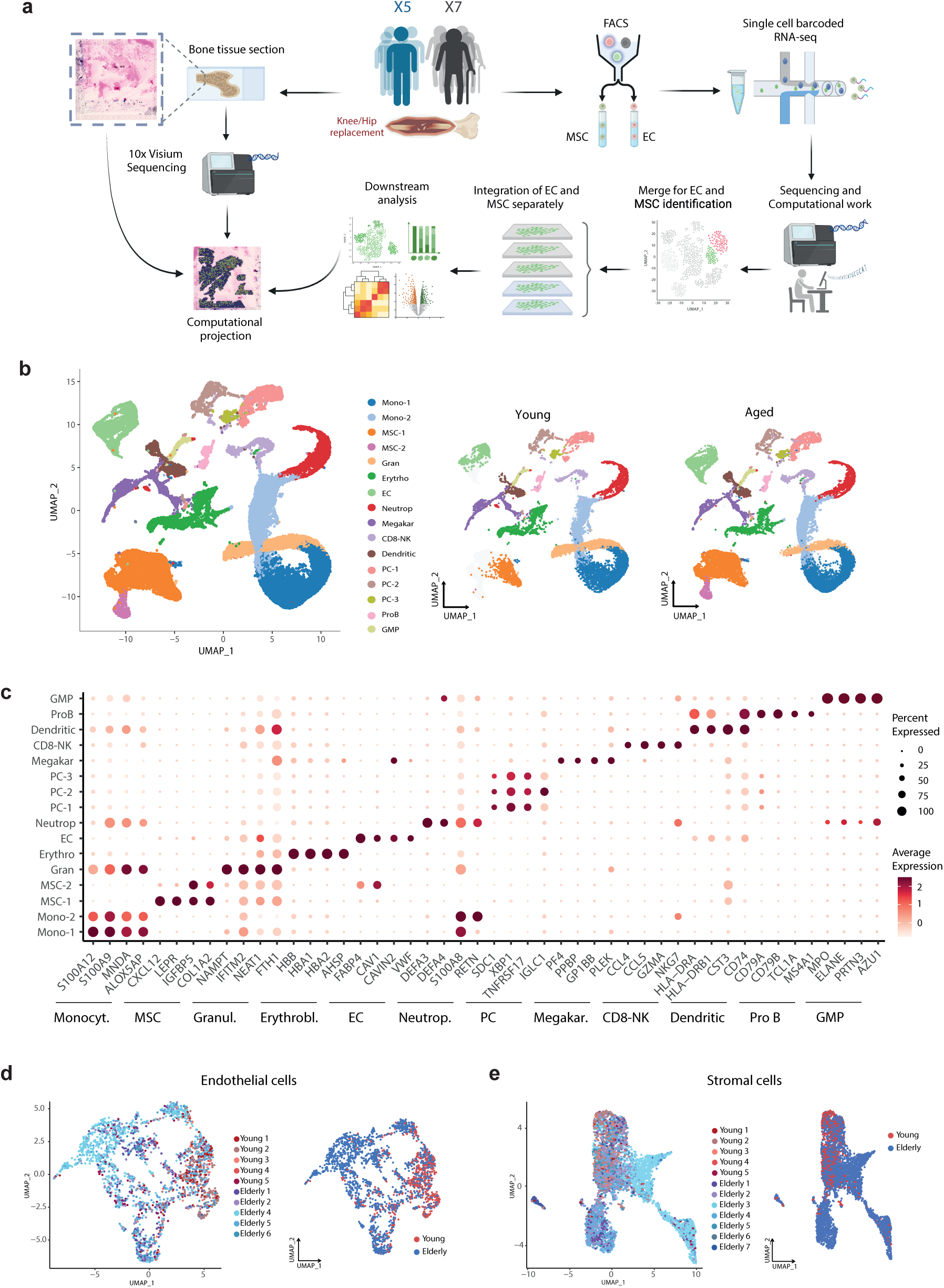
Transcriptional profiling of human young and aged BM endothelial and stromal cells. **a.** Overview of the experimental design for isolation, scRNA-seq, and spatial transcriptomic analysis of EC and MSC from human BM samples. **b.** UMAP representation of human BM microenvironment cells from young and elderly donors, colored by cluster. Right panel: Original UMAP split into young and aged BM datasets. **c**. Dot plot of canonical markers used to define EC, MSC, and hematopoietic populations. The dot size represents the percentage of cells within the cluster that express each gene, and the color indicates the average expression level. **d-e**. Left panel: UMAP projection of the distribution of samples for EC (d) and MSC (e). Right panel: UMAP projection of cells colored by age for EC (d) and MSC (e).

Therefore, we first identified and manually annotated the distinct clusters based on the expression of well-known marker genes (**Fig. 1b, c and Fig. S2a, b**). EC (cluster EC, n=2159 cells) were identified based on the expression of pan-EC markers such as *CD9*, *PECAM1*, *VWF, CDH5,* and *KDR,* and MSC were identified (clusters MSC-1 and MSC-2, n=5317 cells) based on the expression of MSC and osteolineage (OLN)-primed markers including *CXCL12*, *LEPR*, *PDGFRA*, *COL1A1*, and *COL1A2* (**Fig. S2c, d**). The remaining clusters were assigned to well-known hematopoietic BM populations, such as monocytes, granulocyte precursors, erythroblasts, neutrophils, plasma cells (PC), platelets/megakaryocytes, CD8-natural killer-like T cells, dendritic cells (DC), proB cells and granulocyte-monocyte progenitor (GMP), based on the expression of their respective markers (**Fig. 1c**). Notably, the post-annotation resulting cell-type-specific signatures summarized in **Table S2** can serve as a valuable resource for identifying or isolating cell types within BM resident populations, thereby enabling the development of more precise sorting strategies.

To focus on the cells of interest, we separately analyzed EC and MSC from both age groups. Additional filtering and validation were performed to confirm the identity of EC and MSC based on canonical marker expression (**Fig. S2e-f**). UMAPs reveal heterogeneous expression of CD9, CD31, and VWF, providing additional confirmation that the isolated cells originated from bone marrow tissue and were not contaminant EC from muscle or periosteal tissue (*22*). In total, our dataset comprised 2163 EC (549 young, 1614 aged cells) and 5470 stromal cells (808 young, 4662 aged cells) (**Fig. 1d-e and Fig. S2g-h**). Overall, these results demonstrate the feasibility of using BM tissue derived from orthopedic surgeries to successfully isolate and analyze the transcriptional signatures of low-frequency BM EC and MSC in humans.

### Aging is associated with a pro-inflammatory program in EC and the emergence of RAB13^+^ cells

To uncover transcriptional differences between EC from young and aged individuals, we first distinguished the major endothelial vascular beds based on the expression of known molecular markers and gene signatures described previously (*18*, *21*, *23*, *24*) (**Fig. 2a and Fig. S3a**). Arterial cells were characterized by the expression of *PODXL1*, *ICAM2*, *CXCL12, CD34*, *GJA4,* and *HEY1*, whereas sinusoids, overexpressed *STAB2*, *CLEC4G*, *FLT4*, *ENG*, *FNC2*, *VCAM1*, and *LYVE1* (**Fig. 2a**). Beyond characterizing arteries and sinusoids, we identified previously undescribed vascular beds in the human BM, including capillaries and venous vessels (**Fig. 2b, Table S3**). Capillaries were defined by the significant upregulation of known markers, including *RGCC*, *GPIHBP1*, *CA4,* and *EFNB1* (**Fig. 2a**). The venous cluster showed markers of activated post-capillary venules such as *ACKR1*, *POSTN*, *SELE*, and *C7*, alongside higher expression of other genes like *VWF* involved in coagulation and hemostasis(*25*) (**Fig. 2a**). Additionally, we observed a group of EC in a proliferative state (**Fig. S3b**), along with an arterial-like EC subset marked by the expression of *RAB13,* which we designated as RAB13*^+^* cells (**Fig. S3c**). The functional specialization of each EC cluster was further characterized using *Gene Ontology* (GO) *biological process*-based overrepresentation analysis (ORA)(*26*) (**Fig. S3d).**

**Figure 2.**
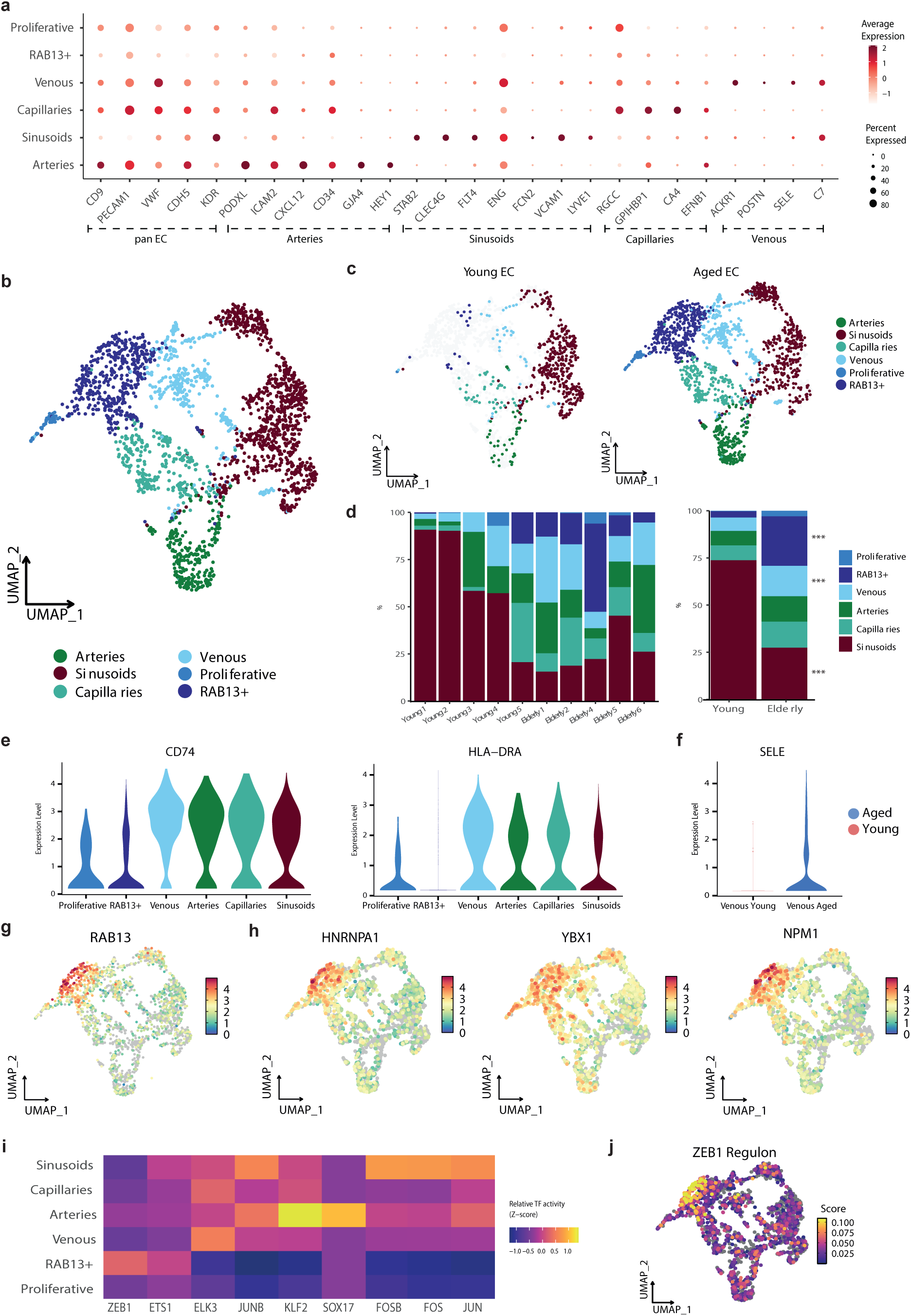
Aging is associated with a pro-inflammatory program in EC and the emergence of RAB13^+^ cells. **a**. Dot plot of markers for EC, arteries, sinusoids, capillaries, and venous used to define EC vascular beds **b**. UMAP depicting vascular subtypes identified within the human BM endothelial compartment. **c**. UMAP visualization of the vascular beds distribution in young (left) and aged (right) EC. **d**. Stacked bar plots showing the proportions of the different vascular states in young and aged EC per sample (left) and age group (right). Asterisks indicate significant differences in the cell proportions between the two groups (FDR < 0.05). **e.** Violin plots showing the expression of genes upregulated in the venous cluster. **f.** Violin plot displaying the expression of *SELE* in young and aged venous EC. **g**. UMAP plot revealing the gene expression of *RAB13* within EC. **h.** UMAP plots showing the expression of genes highly expressed in RAB13^+^ cells. **i**. Heatmap representing z-scaled mean regulon activity score of inferred top-specific regulons for each EC vascular state. **j**. UMAP plot showing the activity score of ZEB1 regulon in EC.

Given the diversity of vascular beds, we first examined how aging affects the composition of EC subtypes in human BM. To estimate robustly the statistical significance, we conducted a permutation-based analysis, and to account for inter-individual variability, we employed a bootstrapping to robustly estimate confidence intervals. Our analysis revealed age-dependent shifts in EC cluster distribution (**Fig. 2c and Fig. S4a)**. Specifically, we observed a significant reduction in sinusoidal EC and an increase in the venous and the RAB13^+^ EC subset in aged individuals (**Fig. 2d and Fig. S4b**). The venous cluster was defined by an increased expression of genes related to “antigen processing and presentation” and “leukocyte activation involved in immune response” (*CD74*, *HLA−DRA*, *HLA−DQB1*, *TFF3*), suggesting an imbalanced immune function and elevated pro-inflammatory response during aging (**Fig. 2e and Fig. S4c**). Besides being a marker of the venous cluster, *SELE*, one of the major drivers of prothrombotic endothelial activation and leukocyte extravasation during inflammation(*27*), was highly expressed in aged venous EC and may serve as a potential biomarker for EC aging (**Fig. 2f and Fig. S4d**).

Interestingly, our study identified the emergence of a transcriptionally distinct subset of EC in aged individuals, defined by high expression of *RAB13*, a GTPase associated with cellular senescence(*28*) (**Fig. 2g, Fig. S4e**). These RAB13*^+^* EC exhibited transcriptional similarities to arterial cells but could be distinguished by elevated expression of genes associated with mRNA stability and translation, including *HNRNPA1*, *YBX1*, and *NPM1* (**Fig. 2h and Table S4**). Functional enrichment analysis of RAB13*^+^* cells revealed an upregulation of ribosomal biogenesis, characterized by increased expression of ribosomal binding proteins (RBP) and translation-related processes (**Table S4 and Fig. S4f, g**). These findings align with recent studies implicating ribosomes in the aging process(*29*). In addition, we observed gene sets associated with Myc targets (*NPM1, HNRNPA1, NAP1L1,* and *EEF1B2*) and cellular senescence (*RAB13, VIM*, *SPARC*, *CCND1,* and *NME2)* in RAB13^+^ cells (**Fig. S4h**).

To better understand the molecular mechanisms underlying this age-related subset, we inferred the active regulons in the RAB13^+^ arterial-like EC population using SCENIC(*30*) and further bioinformatically validated them in SimiC(*31*). Interestingly, our analysis identified the ZEB1 regulon as a candidate regulator of RAB13*^+^* EC subtype (**Fig. 2i, j and Fig. S4i**). The ZEB1 transcription factor (TF) is a known inducer of epithelial-mesenchymal transition (EMT) and has been identified as a critical regulator of HSC functionality and differentiation(*32*). Moreover, recent evidence suggests that Zeb1 induces quiescence of endothelial progenitors during the establishment of vascular homeostasis(*33*).

Altogether, our results identified an elevated inflammatory and prothrombotic response and the emergence of an aging-related EC subset associated with ribosomes and translation elongation, highlighting the remodeling of the human BM vasculature during aging.

### Aged sinusoids exhibited mitochondrial dysfunction and impaired vascular functions

In addition to examining changes in the composition of EC subtypes with aging, we aimed to investigate the transcriptional alterations in the human BM endothelium associated with aging (**Table S5)**. Notably, sinusoidal EC not only decreased in abundance with age but also exhibited the highest number of differentially expressed genes (DEGs), suggesting a pronounced age-related dysfunction in this vascular subtype (**Fig. 3a and Fig. S5a**). Differential expression analysis identified 88 upregulated and 55 downregulated genes in aged sinusoids (**Fig. S5b**). EC exhibited age-related changes that mirror alterations observed in other tissues. Notably, several of these changes contributed to a procoagulant state, thereby increasing the risk of thrombosis in the aging population(*34*). The upregulation of genes such as *ANXA2*, *VWF*, and *PLAT,* among others, in aged sinusoids further supported the hypothesis that age-related endothelial dysfunction may contribute to the heightened incidence of thrombosis (**Fig. 3b and Fig. S5c**). In addition to prothrombotic genes, we also observed elevated expression of *VIM, GSN*, and *ANXA5* (**Fig. 3b and Fig. S5c**). These genes are crucial in ECM rearrangement by modulating cytoskeletal dynamics and cell-ECM interactions, especially in pathologies like cancer invasion, fibrosis, and tissue remodeling. Our findings align with previous studies that identify these genes as potential biomarkers of aging across various cell types and tissues(*35*, *36*). Additionally, aged sinusoids showed upregulation of several genes involved in mitochondrial function and cellular detoxification, such as *ATP5MC2*, *ATP5F1E*, *UQCR11*, *COX4I1*, *COX6A1*, *COX7C*, *NDUFB11,* and *NDUFA11*, reflecting enhanced aging-related dysfunction in cellular respiration (**Fig. 3c, Fig. S5d and Table S6)**. Pathway enrichment analysis further revealed that many of these upregulated genes were associated with key hallmarks of aging, including oxidative stress and adipogenesis (**Fig. S5e**).

**Figure 3:**
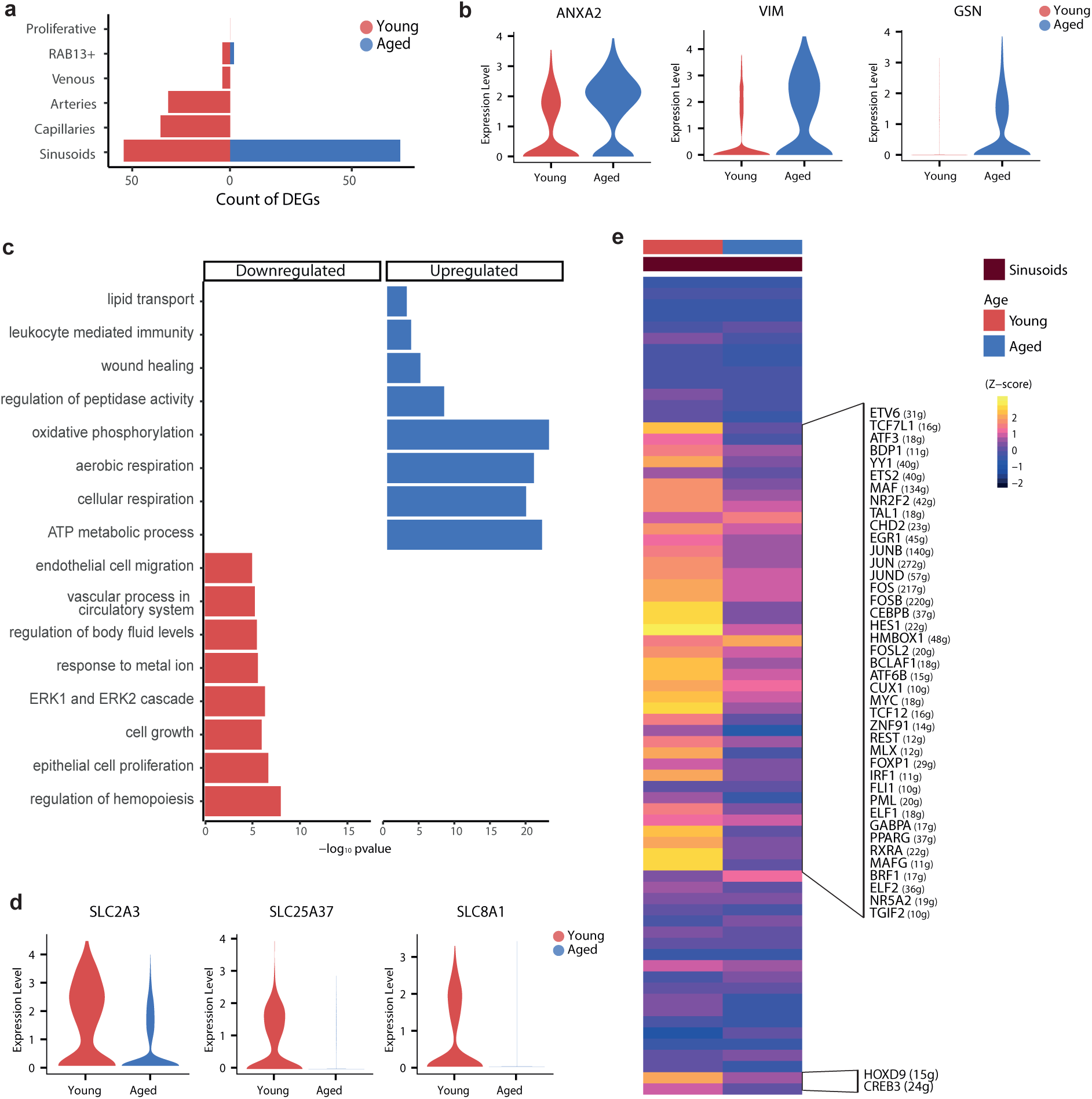
Aged sinusoids exhibited mitochondrial dysfunction and impaired vascular functions. **a**. Plot showing the number of age-related DEGs detected across vascular cell states. **b.** Violin plots of expression of *ANXA2*, *VIM,* and *GSN* in young and aged sinusoids. **c**. Bar charts of enriched GO terms from ORA (p-value < 0.05) comparing DEGs within young and aged sinusoids. The horizontal axis represents the -log10 of p-values. **d**. Violin plots of expression of downregulated solute carriers in aged sinusoids. **e**. Heatmap of mean regulon activity score of significantly enriched regulons in each age group within sinusoids. The color scale represents the z-score of the activity scaled by row.

In contrast, genes involved in endothelial cell proliferation and the regulation of EC migration were downregulated in aged sinusoids, which may be related to the increased vascular permeability and leakiness observed in the aged BM(*37*) (**Fig. 3c and Fig. S5f**). Interestingly, we also observed downregulation of some solute carriers (*SLC2A3*, *SLC25A37*, *SLC8A1*), supporting a decreased response to metal ions (**Fig. 3d**). This decrease also suggested a diminished capacity of EC to process iron and calcium, potentially contributing to the accumulation of non-organic compounds observed during aging, ultimately leading to cellular damage(*38*).

Regulon-based Gene Regulatory Network (GRN) analysis – a transcription factor centered network linking regulators to targets – revealed a global decline in regulon activity in sinusoids along the aging process (**Fig. 3e**). This decline involves Transcription Factors (TF) such as *CEBPB*, *HES1*, *HMBOX1*, *TGIF2,* and *NR5A2*, which are implicated in various biological processes, including cellular proliferation, vascular development, and ion regulation, among others. Collectively, these findings revealed aging-associated molecular features in human BM EC, highlighting thrombosis, mitochondrial dysfunction, compromised vascular functions, and disrupted solute transport as prominent and potential hallmarks in aged sinusoids.

### THY1+-Fibro MSC and matrix remodeling are key factors during BM stromal aging

To explore the age-related transcriptional remodeling of the human BM stroma, we first sought to identify well-established MSC populations in our data based on the expression of established stromal markers(*18*, *21*) (**Fig. 4a and Table S7**). Undifferentiated MSC were defined by the expression of *CXCL12*, *LEPR*, *VCAM1,* and *CP* (**Fig. 4b**). MSC already committed to OLN exhibited elevated expression of osteogenic markers, including *BGLAP*, *SPP1*, *RUNX2,* and *NCAM1*, while the expression of *RGS5*, *MCAM*, *NES*, and *FABP4* distinguished pericytes (**Fig. 4c, d**). We also identified a population of stromal cells highly expressing *LGALS1*, *THY1*, *TIMP1* and *SPARC* that resembles the Fibro and THY1^+^ MSC subsets recently found in a human BM (*21*) (**Fig. S6a, b**). Consequently, we termed this population THY1^+^Fibro-MSC. Despite not capturing a distinctive subset of APOD^+^GSN^high^ MSC as described by Bandyopadhyay et al (*21*), we did observe rare APOD-expressing cells (**Fig. S6c**). In addition, within the undifferentiated MSC, we also identified two clusters expressing *APOB, PPARG*, *CES1, CEBPA,* and *LPL* that may resemble the previously defined adipo-lineage subset (*21*) (**Fig. S6d, e**).

**Figure 4.**
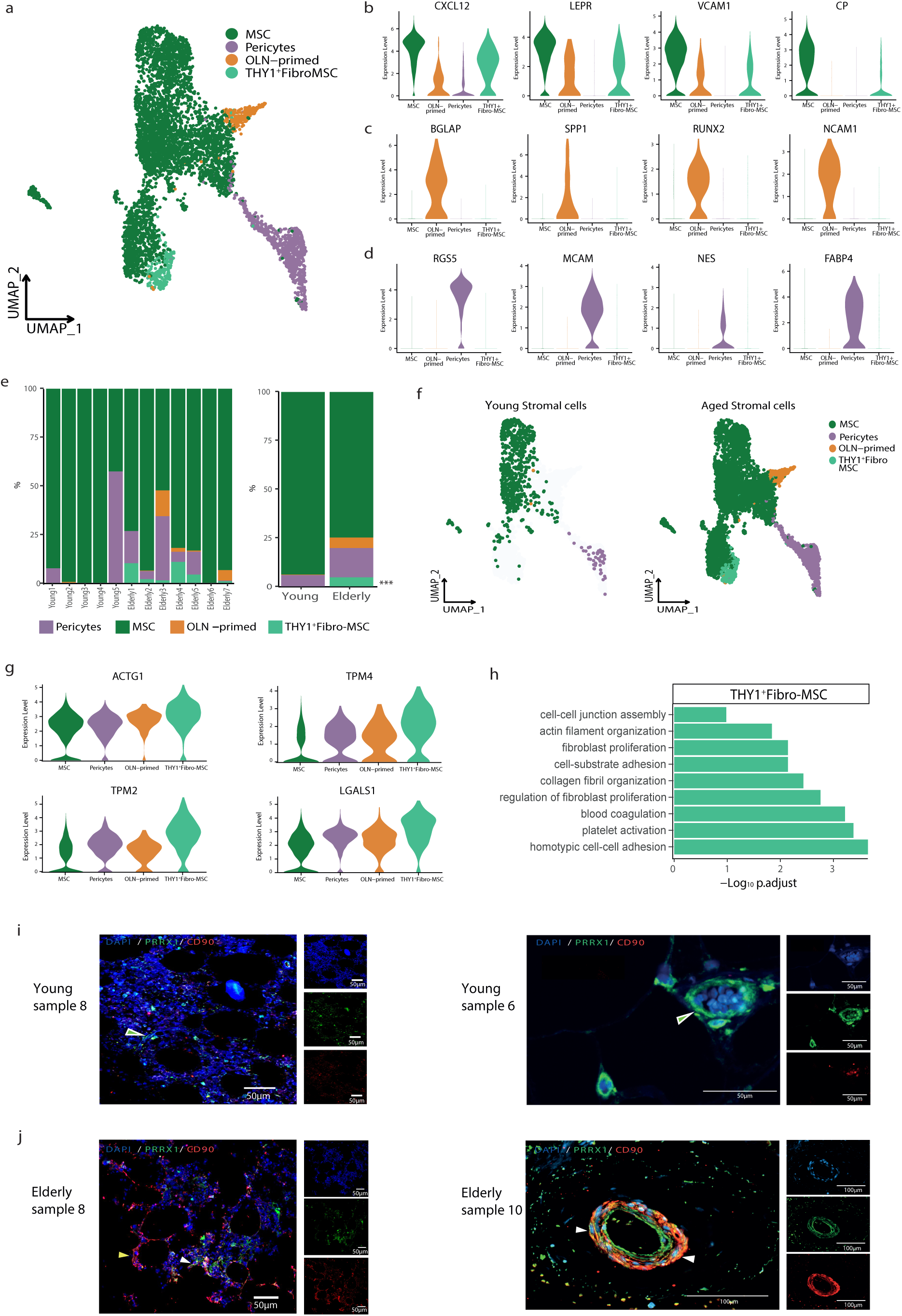
THY1+-Fibro MSC and matrix remodeling are key factors during BM stromal aging. **a**. UMAP projection of the stromal cell subtypes within the human BM mesenchymal compartment. **b-d**. Violin plots showing the expression of well-known markers for MSC (b), OLN-primed (c), and pericytes (d). **e**. Stacked bar plots showing the proportions of the different stromal subtypes states in young and aged MSC per sample (left) and age group (right). Asterisks indicate significant differences in the cell proportions between the two groups. **f**. UMAP visualization of the stromal subtypes in young (left) and aged (right) MSC. **g**. Violin plot showing the expression of matrix-related genes upregulated in the THY1+Fibro-MSC cluster. **h**. Bar chart of enriched GO terms from ORA (p-value < 0.05) defining the THY1+Fibro-MSC cluster. The horizontal axis represents the -log10 of adjusted p-values. **i-j**. Immunofluorescence (IF) staining of THY1^+^ Fibro stromal cells (CD90^+^) (red), MSC (PRRX1^+^) (green), and nucleus (DAPI) (blue) in FFPE biopsies **(Table S9)** from two young (i) and two elderly (j) individuals. Scale bars: 100 μm. Green arrows indicate MSC PRRX1+, white arrows indicate MSC THY1+ (PRRX1+ THY1+), yellow arrows indicate THY1+ cells.

To assess the impact of aging on the stromal compartment at the cellular level, we investigated potential changes in the abundance of each cell type between young and aged MSC. Our analysis revealed significant variation in cellular composition among individuals (**Fig. 4e, f, and Fig. S6f**). Notably, the presence of pericytes and OLN-primed cells was observed only in specific samples, which may not accurately reflect age-related changes in proportions, possibly due to variations in sample dissociation. Nevertheless, we consistently found THY1^+^Fibro-MSC in all samples from elderly individuals, suggesting that these cells emerge in the aged BM (**Fig. 4e left panel**). The expansion of these THY1^+^Fibro-MSC during aging aligns with the identification of this mesenchymal subset by Bandyopadhyay et al (*21*), whose cohort comprises elderly individuals, thereby approximating a relatively “aged” BM microenvironment.

To delve into the potential role of THY1^+^Fibro-MSC as regulators of BM decline during the aging process, we analyzed their transcriptome (**Table S8)**. We observed increased expression of *FN1*, *MIF*, *S100A6,* and *ANXA2,* suggesting the profibrotic nature of aged MSC (**Fig. S6g)**. Besides the expression of multiple genes implicated in ECM organization and cell-matrix interactions, such as *ACTG1*, *TPM2*, *TPM4*, and *LGALS1* (**Fig. 4g**), THY1^+^Fibro-MSC showed elevated expression of *COL1A1*, *COL3A1,* and *COL1A2*, which are involved in the TGF-β signaling pathway (**Fig. S6h**). Interestingly, TGF-β signaling plays a pivotal role in myelofibrosis by promoting BM fibrosis and collagen deposition(*39*). Moreover, these cells exhibited high expression of key genes implicated in myelofibrosis and other fibrotic disorders, such as *IGFBP6* and *IGFBP7* (**Fig. S6i**)(*40*). In line with these results, pathway analysis identified significant enrichment in fibroblast proliferation, matrix, and EMT, suggesting the implication of this profibrotic state in aged MSC (**Fig. 4h, Fig. S6j and Table S8**). THY1^+^Fibro-MSC also presented enrichment in homophilic cell adhesion, suggesting an altered interaction with the marrow environment (**Fig. 4h)**.

To further confirm the changes in MSC between young and elderly BM samples, including the identification of THY1^+^Fibro stromal cells in aged individuals, we performed immunofluorescence (IF) staining of this population in new formalin-fixed paraffin-embedded (FFPE) young and elderly BM biopsies (**Table S9**). An increase in THY1^+^ cells among aged MSC, identified by expression of CD90^+^ in PRRX1^+^ cells, was observed (**Fig. 4i, j and Fig. S6k**). In brief, these results suggest a more fibrotic and less functional aged BM microenvironment, potentially contributing to the reduced hematopoietic-supportive capacity in aged BM.

Building on these findings, we hypothesize that THY1^+^ MSC may contribute to the age-dependent development of fibrosis and may have important implications for BM fibrotic diseases such as myelofibrosis. Using IF staining, we observed an increase in THY1^+^ MSC, marked by an increase in the number of cells co-expressing CD90 and PRRX1 in the BM of myelofibrosis patients, particularly in scar tissue areas (**Fig. S6k-m and Table S9**). Taken together, the elevated presence of these profibrotic cells in myelofibrosis patients indicates their potential involvement in the replacement of hematopoietic cells with fibrotic tissue. Although functional validation of this hypothesis is necessary, our findings suggest that these cells could be key in the age-related deterioration of the BM microenvironment and hematopoiesis.

### Impaired differentiation and oxidative metabolism in aged MSC

To further delve into the different molecular processes underlying stromal aging, we investigated the transcriptional changes between young and aged stromal cells (**Table S10**). Undifferentiated MSC showed the largest number of DEGs, demonstrating age-related defects in these mesenchymal progenitors (**Fig. 5a and Fig. S7a**). Specifically, differential expression analysis detected 367 DEGs, including 203 upregulated genes and 164 downregulated genes in aged MSC (**Fig. 5b**). Enrichment analysis revealed that many downregulated genes were linked to ossification-related processes, suggesting that age-related bone loss may stem from impairments in the early stages of differentiation (**Fig. 5c and Table S11**). We also found a downregulation of genes that regulate hematopoiesis and cell differentiation, such as *MEIS2*, *GAS6*, *SMAD7,* and *ID2*, as well as heat-shock proteins and other essential genes mediating the response to protein folding and external stimulus (*HSPA1A*, *HSPA1B*, *NEAT1* and *THBS1* among others) (**Fig. S7b, c)**. Moreover, consistent with previous findings in other tissues, the downregulation of angiogenesis-related terms in aged BM MSC suggests a compromised ability of the mesenchymal progenitors to support BM vascular networks(*41*).

**Figure 5.**
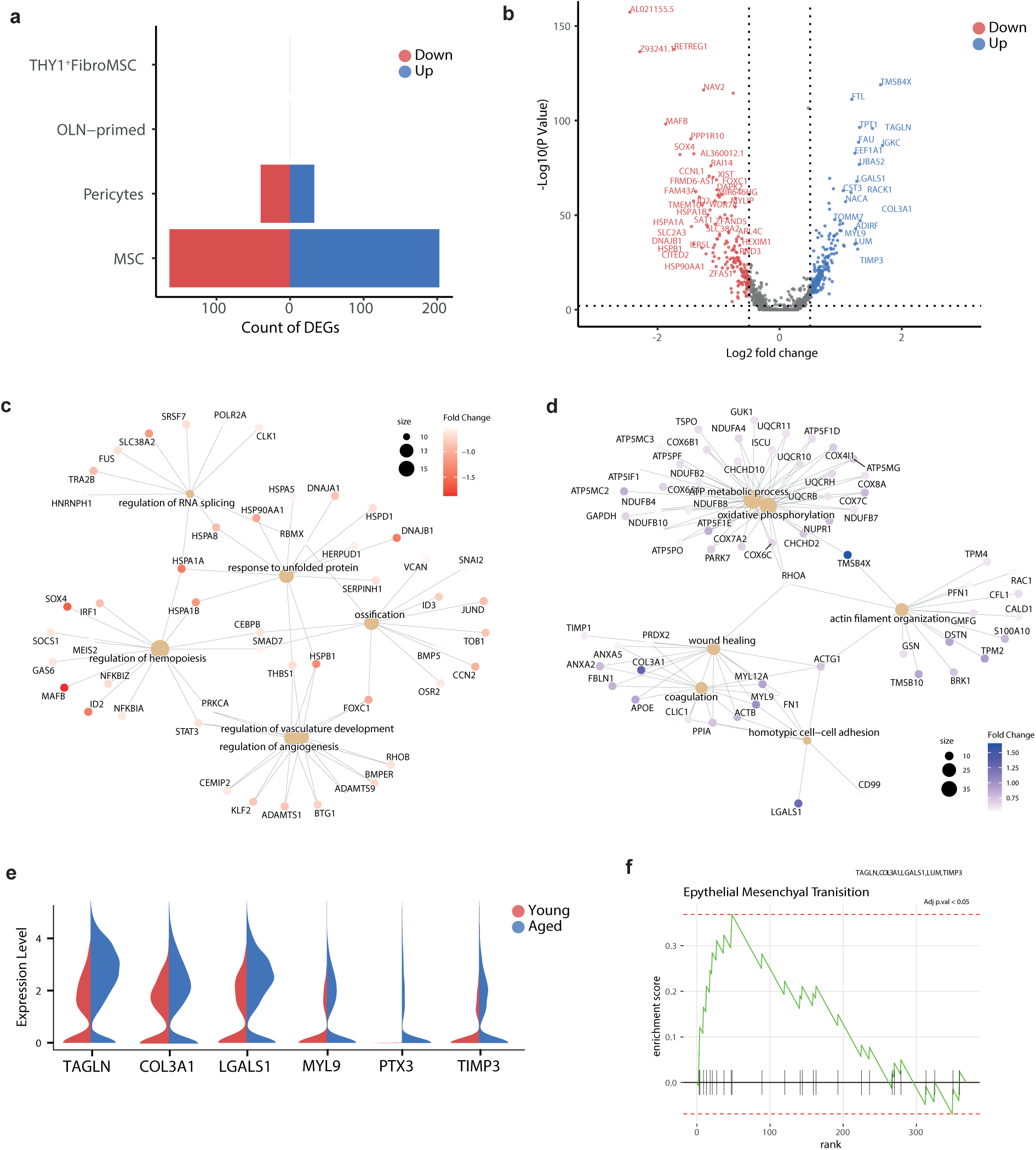
Impaired differentiation and oxidative metabolism in aged MSC. **a**. Plot showing the number of age-related DEGs detected across the BM stromal subpopulation. **b**. Volcano plot of the DEGs between young and aged MSC. The y-axis represents the -log10(p-value), and the x-axis represents the log2(fold change). The color of the dot denotes the age group for which DEG was detected, with grey dots representing non-significant genes. **c**. Cnetplot showing the links between genes and biological processes downregulated in aged MSC. Node size reflects the number of significantly enriched genes in the node and colors the log_2_ Fold change expression of each gene. **d**. Cnetplot showing the relationship among individual GO terms and genes upregulated in aged MSC. Node size indicates the number of significantly enriched genes in the node and colors the log_2_ Fold change expression of each gene. **e**. Split violin plots showing the expression of aging-related matrix remodeling-associated genes upregulated in aged MSC. **f**. GSEA plot of “Epithelial Mesenchymal Transition” term significantly enriched in aged MSC.

Additionally, aged MSC showed increased expression of genes associated with OXPHOS and ATP metabolism, including *GSTP1*, *ATP5F1E*, *COX7C*, *NUPR1*, *NDUFA4,* and *CHCHD2*, suggesting an abnormal mitochondrial metabolism in aged MSC (**Fig. 5d and Fig. S7d, e**). This could be attributed to mitochondrial dysfunction and elevated ROS levels, in line with the reported impairment of mitochondrial complexes in aged MSC(*42*). The expression of the adipogenic genes *APOE*, *SPARCL1,* and *CAVIN1,* and several genes involved in ECM organization and remodeling, such as *TAGLN, COL3A1, LGALS1*, *MYL9*, *PTX3,* and *TIMP3*, was increased with age (**Table S10** and **Fig. 5e**). Accordingly, Gene Set Enrichment Analysis (GSEA) revealed enrichment in EMT, OXPHOS, and adipogenesis (**Fig. 5f and Fig. S7f**). Along with the enhanced EMT process during aging, aged MSC showed decreased expression of metalloproteinases *ADAMTS9* and *ADAMTSL3,* further indicating the role of the ECM remodeling during BM stromal aging (**Fig. S7g**). This is aligned with the fact that MSC senescence induces changes in ECM, affecting their repair tissue ability(*43*). In summary, our results suggest the contribution of MSC to the age-related alterations in hematopoiesis. The observation that MSC from aged subjects show an increased oxidative metabolism, EMT, impaired hematopoietic differentiation, and protein folding capacity, along with the presence of THY1^+^Fibro-MSC, supports this hypothesis and highlights the BM niche as a target for anti-aging interventions.

### Age-related remodeling of EC-MSC communication

After characterizing the aging-associated changes in the non-hematopoietic BM microenvironment, we aimed to dissect how these alterations may affect cell-cell communication mechanisms. To this end, we conducted a single-cell-based ligand-receptor (L-R) analysis using LIANA(*44*), focusing initially on age-related changes in the interactions between EC and MSC (**Fig. 6a, Table S12**).

**Figure 6.**
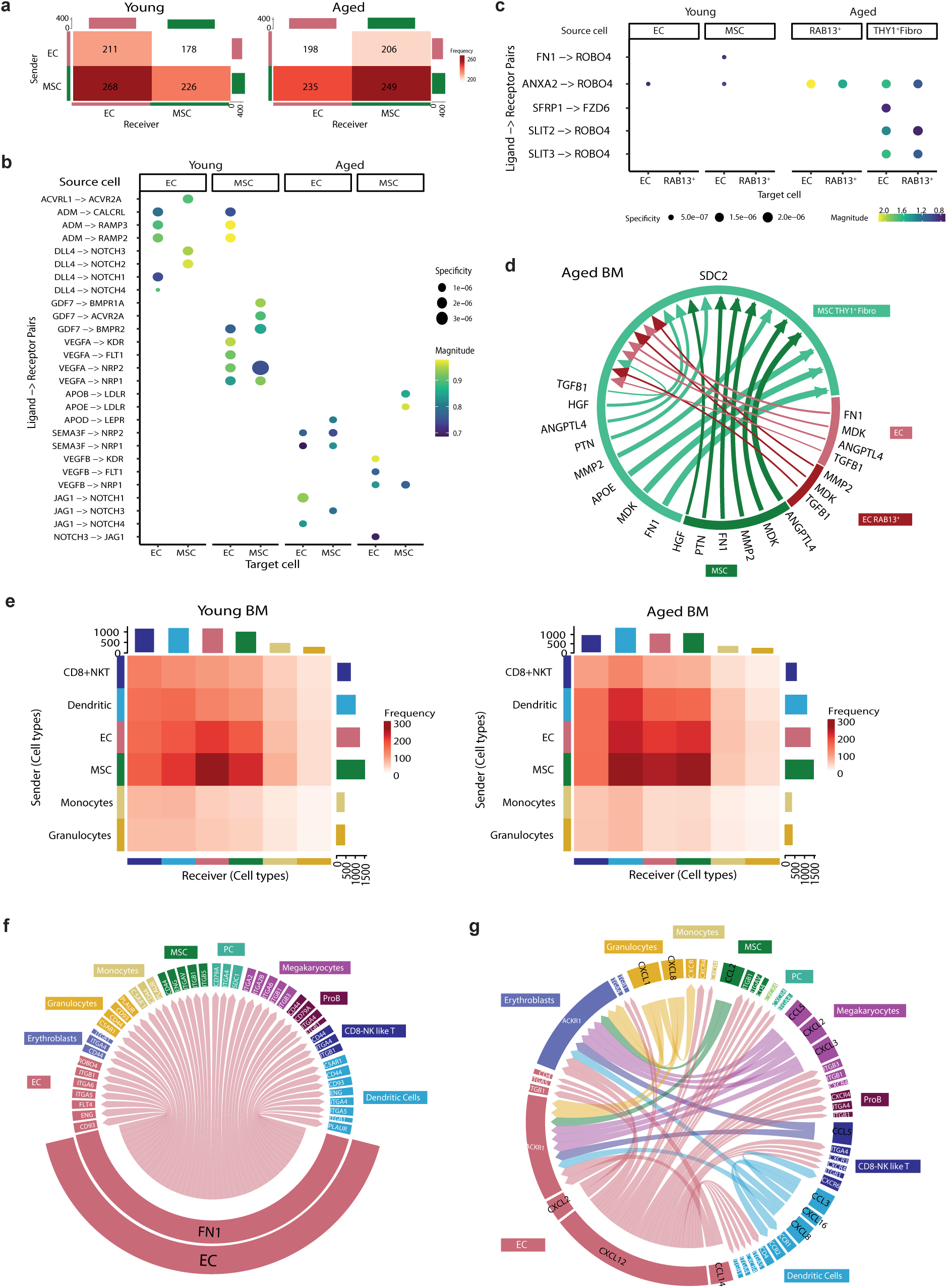
Age-related remodeling of EC-MSC communication and the BM interactome. **a**. Heatmap illustrates predicted cellular communication between EC and MSC in young and aged BM. **b**. Dot plots representing specific L-R interactions between EC and MSC in young and elderly donors. Dot size and color indicate the specificity and the strength of the interaction, respectively. **c**. Dot plots representing L-R communication pairs involving the RAB13^+^ EC state and THY1^+^Fibro-MSC population in young and aged BM. Dot size and color indicate the specificity and the strength of the interaction, respectively. **d**. Chord diagram depicting age-specific communication of THY1^+^Fibro-MSC population through *SDC2* receptor. Color represents the signal sender, and width is the strength of interactions. **e**. Heatmap illustrates predicted cellular interactions between BM microenvironment cells (EC, MSC, CD8+ NKT cells, dendritic cells, monocytes, and granulocytes) in the young (left) and aged (right) BM. Block sizes and red coloration are proportional to the number of L-R pairs. **f**. Chord diagram depicting age-specific interactions between *FN1* from EC and several proteins from different cell types. Color represents the signal sender, and width is the strength of interactions. **g**. Chord diagram showing the significant age-related signaling network between CXCL and CCL family members and other proteins from various cell types. Colors and widths represent the signal senders and the strength of interactions, respectively.

The remodeling of EC and MSC interactions supported the vascular weakening observed in the BM with aging. Specifically, *DLL4*, a Notch ligand crucial for angiogenesis(*45*), interacted with *NOTCH1-4* in young EC and MSC. In contrast, in aged EC, *DLL4* interactions were replaced by *JAG1,* suggesting an impairment in blood vessel formation (**Fig. 6b**). This observation was further supported by age-related vascular endothelial growth factor (VEGF) family alterations. In young BM, *VEGFA* mediated interactions with endothelial receptors *KDR*, *FLT1, NRP1,* and *NRP2*, whereas *VEGFB* was predominant in aged individuals (**Fig. 6b**). Additionally, interactions mediated by *GDF7* and bone morphogenetic protein (BMP) elements, presented in the young stroma, were notably absent in aged cells (**Fig. 6b**). In contrast, the aged BM microenvironment showed a more adipogenic phenotype as evidenced by the interactions involving apolipoproteins (*APOB* and *APOE* with *LDLR* in MSC and *APOD* from EC with *LEPR* in MSC) (**Fig. 6b**). These changes in EC-MSC communication in the aged BM is consistent with the observed shift in MSC differentiation towards adipocytes at the expense of osteoblast differentiation.

A more detailed analysis of the interactions between the RAB13^+^ arterial-like EC and the THY1^+^Fibro-MSC populations identified specific L–R pairs that became more prominent in the aged BM (**Fig. 6c**). Interactions like *SLIT2*-*ROBO4*, *SLIT3*-*ROBO4,* and *SFRP1*-*FZD6* were only identified in aged individuals. The expression of *SRFP1* and *FZD6* has been described as increasing in aged senescent cells, suggesting their potential as markers of Wnt signaling and senescence-related aging in the BM(*46*, *47*). Additionally, aged THY1^+^Fibro-MSC exhibited extensive crosstalk with both EC and MSC subpopulations through *SDC2* receptor interactions, implicating cell adhesion and ECM processes in the aging BM microenvironment (**Fig. 6d**). Overall, our findings highlight key L-R interactions that may contribute to the rewiring of aged EC and MSC, uncovering key molecular pathways involved in BM aging.

### Impact of aging on the BM microenvironment interactome

Next, we inferred the cell-cell interactions between EC, MSC, and the broader BM microenvironment during aging (**Fig. S8a-b**). Interestingly, interactions with DC increased with age, possibly influenced by the chronic inflammation commonly observed in elderly individuals(*48*) (**Fig. 6e**). By examining the age-related changes on specific ligands and receptors, *FN1* appeared as a putative interactor of the aged BM microenvironment (**Fig. 6f**). *FN1* ligand express by MSC, DC, and mainly EC interacts with different members of the integrins family further implicating ECM remodeling in BM aging. Furthermore, predicted *TGFB1*-mediated interactions increased in aged EC and MSC compared to the young BM (**Fig. S8c**). The interactions of *TGFB1* with its receptors and several integrins, such as *ITGAV* on MSC, which were specific to the aged BM microenvironment, suggest a possible contribution to age-related fibrosis.

The elevated inflammatory cytokine signaling aligns with the pro-inflammatory state described in the aged BM (**Fig. 6g**). *CXCL12* from aged EC interacted with EC, MSC, and DC through *CD4.* Additionally, *ACKR1* from EC and erythroblasts interacted with various hematopoietic cell types via CXCL family members (*CXCL8*, *CXCL2*, *CXCL1*) and CCL chemokines (*CCL14*). This shift toward a pro-inflammatory state during aging could be accentuated by the increased interactions mediated through *MIF*, *CD74*, and *CD99* ligands observed in the aged BM microenvironment (**Fig. S8d**).

In conclusion, the remodeling of the BM interactome during aging suggests that non-hematopoietic components, particularly through increased communication with DC, may contribute to the pro-inflammatory microenvironment observed in aged individuals. Additionally, the *FN1* and *TGFB1* pathways warrant further investigation as potential targets for mitigating fibrotic processes in the aging BM.

### Spatial transcriptomics supports age-associated vascular changes

We have characterized the age-related shifts in BM non-hematopoietic cell proportions, molecular profiles, and cell-to-cell communication. To gain additional evidence at spatial resolution of these changes, we employed spatial transcriptomics using FFPE BM from two new samples, one young (34 years of age) and one elderly (76 years of age), using the Visium Gene Expression platform (10X Genomics)(*49*) (**Table S9**). After pre-processing and quality control analysis, we detected 757 and 810 spots in young and elderly samples, respectively, with an average of 612 features and 1417 counts per spot in the young and 561 features and 1415 counts in the elderly (data not shown).

Given our interest in characterizing aging-related shifts in EC and MSC, we first aimed to identify the spots with a higher likelihood of containing these specific cell types. To label the spots, we leveraged a scRNA-seq atlas from Bandyopadhyay et al (*21*), which also include HSC, as a reference for the annotation analysis. First, we estimated both cell proportion and a signature score per cell type per spot (**Fig. S9a-c**). Then, for each cell type, spots highly ranked for both measures were considered “high-confidence spots” for such cell type as detailed in supplemental methods (**Fig. S9d-f**). All analyses were conducted separately for both samples, resulting in a subset of spots that mapped EC, MSC, as well as hematopoietic cells, including HSC, B cells, DC, erythroblasts, monocytes, neutrophils, PC, and T cells in the human BM (**Fig. 7a and data not shown)**. The spatial spots of endothelial and stromal cell populations were confirmed using IF and Hematoxylin-Eosin (H&E) staining (**Fig. 7b and Fig. S9g**). Specifically, non-hematopoietic niche cells were localized around blood vessels and identified by IF using the colocalization of specific markers for EC (CD31^+^ CD45^-^) and MSC (PRRX1^+^ CD45^-^).

**Figure 7.**
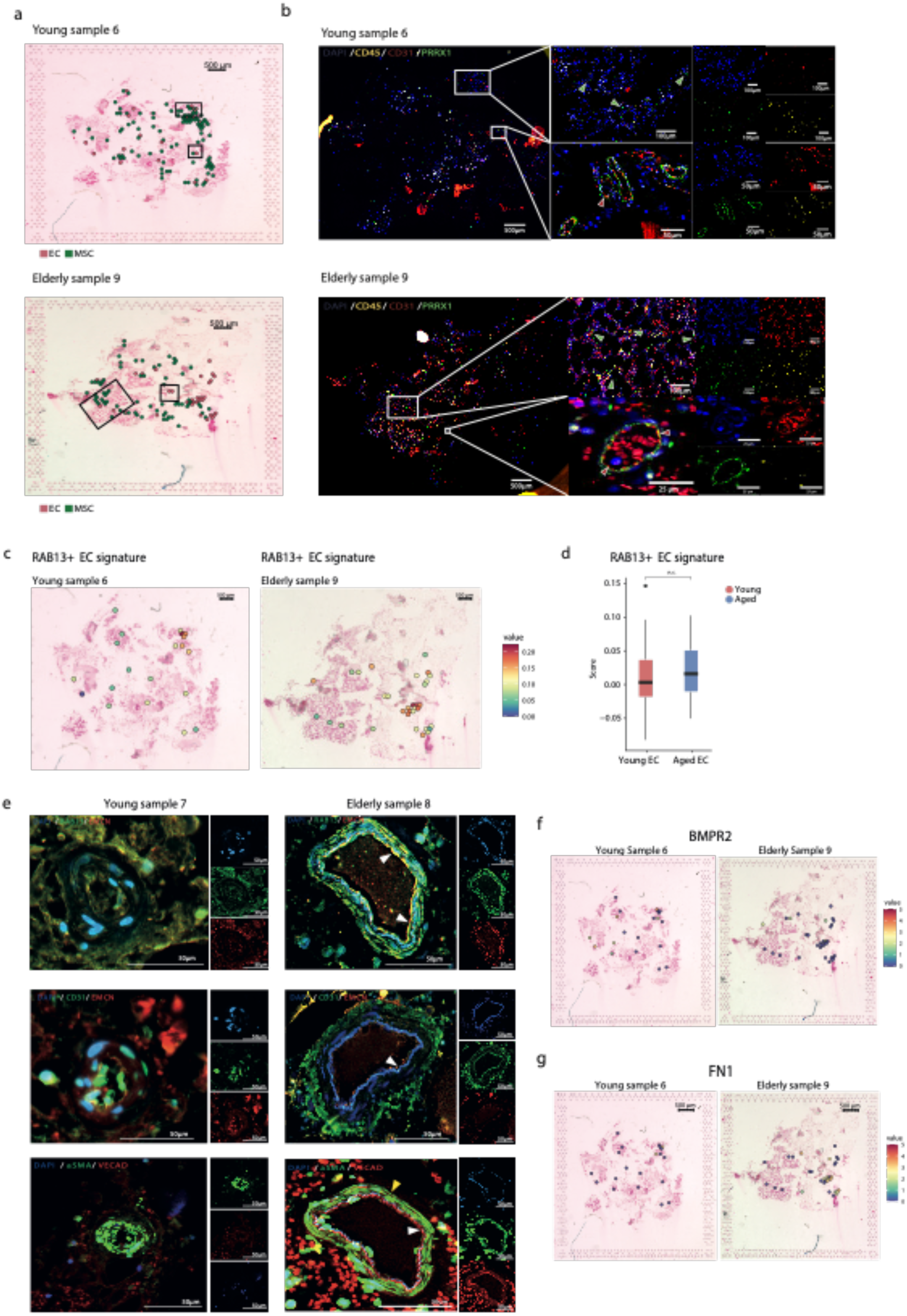
Spatial transcriptomics supports age-associated vascular changes. **a**. Pie charts on H&E staining illustrate the proportion of each cell type highly contributing to the transcriptomic signature of each spot in young sample 6 (upper) and elderly sample 9 (bottom) new BM samples **(Table S9)**. **b**. IF staining of hematopoietic cells (CD45^+^) (yellow), EC (CD31^+^) (red), MSC (PRRX1^+^) (green), and nucleus (DAPI) (blue), in young (upper panels) and elderly (bottom panels) BM samples. Scale bars are indicated on each panel. Red arrows indicate the presence of EC (CD31+), and green arrows indicate the presence of MSC (PRRX1+). **c**. Spatial expression patterns of the signature from RAB13^+^ EC identified in scRNA-seq data in EC-labelled spots of young (left panel) and elderly (right panel) BM samples. **d**. Box plots displaying the RAB13^+^ EC signature in young (red) and elderly (blue) EC. **e**. IF staining in the FFPE BM biopsies **(Table S9)** of the young sample 7 (left) and elderly sample 8 (right). Scale bars: 50 μm. Upper panel: RAB13 (green), EMCN (red), and DAPI (blue). White arrows indicate RAB13 EC (RAB13+ EMCN+). Middle panel: CD31 (green), EMCN (red), and DAPI (blue). White arrows indicate EC (CD31+ EMCN+). Lower panel: Alpha Smooth Muscle Actin (aSMA)(green), VE-Cadherin (VECAD) (red), and DAPI (blue). White arrows indicate EC (VCAD+), and yellow arrow indicates vascular smooth muscle cells (aSMA+). **f**. Spatial expression patterns of the BMPR2 gene in EC golden spots of young (left panels) and elderly (right panels) BM samples. **g.** Spatial expression patterns of the *FN1* ligand in EC golden spots of young (left panels) and elderly (right panels) BM samples.

Next, we examined the spatial expression patterns of age-related transcriptional changes identified in our scRNA-seq analysis, with a focus on EC. Even though we lack sufficient statistical power, we observed higher values of the RAB13-related signature in aged EC spots (**Fig. 7c-d**). Moreover, RAB13^+^ EC colocalized with the spatial expression pattern of *YBX1*, a TF involved in RNA splicing and translation control, which was shown to be upregulated in the RAB13^+^ arterial subset identified in our scRNAseq-data (**Fig. S10a)**. The spatial enrichment of this EC subset with aging was also confirmed through IF staining of RAB13^+^ cells from FFPE BM biopsies (**Fig. 7e**). RAB13^+^ EC were more abundant in the aged BM, consistent with our previous transcriptomic findings. It is noteworthy that RAB13 expression was predominantly localized near the periarteriolar regions, as evidenced by the proximity of RAB13^+^ CD31^+^ EC to α-SMA1^+^ cells (**Fig. 7e**).

Consistent with the significant age-related increase in the venous cluster and supporting the “inflammation and immune” hypothesis from the transcriptomic data, we found higher spatial expression of *CLU* and *CD74* in aged EC (**Fig. S10b**). Additionally, aged EC-labeled spots showed a tendency towards oxidative metabolism, which aligns with the mitochondrial dysfunction identified in aged sinusoids through our scRNA-seq analysis (**Fig. S10c**). *VIM* and *GSN*, two genes implicated in ECM rearrangement and previously found upregulated in aged sinusoids, were highly expressed in aged EC (**Fig. S10d**).

Finally, spatial resolution analysis of L-R interactions supported findings from our previous BM microenvironment cell-to-cell communication analysis. Specifically, we observed reduced *BMPR2* expression in spots enriched in aged EC, indicating diminished support for osteogenic differentiation through EC-MSC interactions in the aged BM (**Fig. 7f and Fig. S10e)**. Conversely, *JAG1-NOTCH3* signaling was detected in spots enriched in aged EC (**Fig. S10f)**. In line with the increased interactions between niche cells and DC identified by our scRNA-seq analysis, EC-labeled spots in aged BM showed a positive correlation with the DC score, suggesting a role for EC-DC communication in the pro-inflammatory aged BM (**Fig. S10g)**. Interestingly, we observed a higher spatial expression of the *FN1* ligand in spots enriched in aged EC, supporting our previous transcriptomic findings pointing FN1 as a key interactor in the aged BM (**Fig. 7g and Fig. S10h)**.

Overall, our findings revealed spatially distinct, age-related changes within the human non-hematopoietic BM microenvironment, with an impact on EC. Supported by our scRNA-seq analysis, these results suggest alterations in cell state distribution, signaling pathways, and intercellular communication within the aged BM.

## Discussion

Over the past decades, significant progress has been made in elucidating the mechanisms of human BM aging. Advances in scRNA-seq have provided unprecedented resolution into the heterogeneity and age-related trajectories of HSC(*7*, *50*, *51*). Parallel studies of the aged HSC niche have uncovered widespread alterations in the BM microenvironment that actively contribute to the functional decline of HSC over time (*51–54*). These findings have underscored the BM niche as a promising target for therapeutic rejuvenation strategies. Nonetheless, due to the inherent challenges of studying aging in humans, much of our current knowledge stems from murine models (9, 55,56).

Here, we provide a comprehensive transcriptional characterization of the changes that occur during physiological aging of the BM non-hematopoietic microenvironment with a special focus on the endothelial and mesenchymal compartments. Additionally, using IF staining and spatial transcriptomic profiling, we provide valuable preliminary insights into the spatial organization of these cells. Most significantly, we provide support for some of the findings derived from the scRNA-seq analysis of BM aging.

Our findings expand on the well-characterized arteries and sinusoids, uncovering previously undescribed vascular subtypes within the human BM endothelium, including capillaries and veins. We revealed significant age-dependent shifts in EC composition, with an increased prevalence of the venous cluster in elderly samples, marked by genes related to immune activation and coagulation. This indicates a shift towards enhanced inflammation and elevated thrombotic activity with aging in EC. Interestingly, we uncovered an arterial-related cluster in aged EC, characterized by the high expression of *RAB13* associated with dysregulation of translational elongation activity and ribosomal processes. The spatial enrichment of this arterial-associated EC subset during aging was further demonstrated by IF staining, showing the colocalization of RAB13^+^ cells with CD31^+^ EC and α-SMA1^+^ cells. This newly identified aging-related subset, revealed through scRNA-seq and spatial transcriptomics, is potentially regulated by the TF ZEB1(*33*) and demonstrates elevated expression of RBP, positioning these cells bas candidate regulators of aging and promoters of cellular senescence(*29*, *57*). The role of RAB13 in cancer progression is well-documented, making the identification of RAB13^+^ cells as a novel subset of aged EC a potential therapeutic target(*58*). Sinusoids were identified as the most distinct vascular bed during aging, exhibiting a prothrombotic profile, mitochondrial dysfunction, and a decline in transcriptional activity. Aged sinusoids also showed a signature associated with an impaired ability to maintain vascular integrity and angiogenic potential, possibly promoting regression in vascular structure and disrupting inflammatory homeostasis, consistent with previous studies conducted in mice(*11*, *55*, *59*).

Despite considerable advancements in characterizing the human BM stroma, age-related transcriptional changes in MSC remain unresolved(*19*, *21*, *60*, *61*). Notably, our results revealed the emergence of THY1^+^Fibro-MSC in the aged stroma, which exhibited high similarity to the recently identified THY1^+^ Fibro-MSC subpopulations(*21*). These cells showed an enrichment of genes associated with TGF-β signaling pathway, ECM, and EMT, suggesting the implication of these profibrotic processes in aged MSC. We also confirmed histologically that CD90^+^ BM MSC coexist with fibrosis(*62*). Furthermore, analyzing the transcriptional changes in MSC during aging showed an increase of ATP metabolism and OXPHOS in aged individuals, reduced unfolded protein response (UPR), and impaired support for hematopoietic and osteoblast differentiation. Overall, our findings reveal extensive transcriptional remodeling within the endothelial and mesenchymal compartments of the BM, characterized by increased inflammation, impaired ribosomal and mitochondrial function, and elevated ROS production. These age-specific changes align with established hallmarks of aging(*63*, *64*), positioning EC and MSC as promising targets for strategies aimed at rejuvenating the aged BM microenvironment(*65*, *66*).

The differences in the BM microenvironment significantly influence the behavior and regenerative capacity of aged HSC(*67*). Therefore, studying this crosstalk is crucial for developing therapeutic strategies to mitigate age-related decline in HSC function and enhance health outcomes in older individuals. The shift from *VEGFA* to *VEGFB* interactions and the altered NOTCH signaling observed in aged EC may contribute to impaired blood vessel formation during aging(*68*). This disruption of NOTCH signaling supports previous studies highlighting its role in maintaining HSC stemness(*11*, *69*). Moreover, the aged BM niche showed increased apolipoprotein interactions, consistent with the accumulation of adipocytes during BM aging and the compromised differentiation of aged MSC(*13*). Remarkably, we identified *SFRP1-FZD6* and *SDC2* receptor interactions as potential key regulators of senescence and ECM processes related to the emergence of RAB13^+^ EC state and THY1^+^Fibro-MSC population during aging. *FN1* and *TGFB1* emerged as central signaling hubs in the aged BM microenvironment, reflecting impaired matrix remodeling and potentially driving cell activation and inflammatory processes(*70*). This shift toward a pro-inflammatory state during the aging process was reinforced by the enhanced communication with DC and the increased interactions mediated by cytokine signaling, *MIF*, *CD74*, and *CD99*.

Despite the technical limitations associated with applying spatial transcriptomics to mineralized tissues and BM in particular (*71*, *72*), we were able to perform Visium Gene Expression technology from 10X Genomics on human FFPE BM biopsies from young and elderly individuals. We mapped the spatial distribution of hematopoietic and non-hematopoietic components within the human BM niche, which was supported by IF staining for CD45, CD31, and PRRX1. Our data provided initial spatial context for the EC changes previously observed in our scRNA-seq analysis. Notably, we observed an increased spatial localization of RAB13^+^ senescent cells near periarteriolar regions, alongside distal EC showing elevated expression of *GSN* in the aged BM. Additionally, our spatial transcriptomic data supported the enhanced communication between aged EC and DC, and the FN1 signaling pathway as a potential interaction hub in the aged BM.

In summary, our study provides novel insights into the age-related remodeling of the EC and MSC transcriptome, interactome, and spatial architecture, offering potential explanations for the deterioration of the hematopoietic system observed during aging. We find that aging compromises the ability of EC to maintain vascular integrity and reduces the protein folding capacity and support for hematopoietic differentiation of MSC. Both cell types exhibit compromised mitochondrial function and heightened inflammatory and prothrombotic properties with age. We also identified an age-related EC subset responsible for translation and ribosomal processes. Moreover, we observe a more fibrotic and less functional aged BM stroma influenced by THY1^+^Fibro-MSC, which may contribute to its diminished capacity to support hematopoiesis in aging-related pathologies. Our spatial transcriptomics analysis supports our previous scRNA-seq results, highlighting the presence of RAB13^+^ senescent cells in aged EC and age-associated interactions. Collectively, our results highlight the crucial role of non-hematopoietic components in shaping the functional trajectory and longevity of HSC. These age-related alterations position the non-hematopoietic BM niche as a promising therapeutic target for mitigating hematopoietic decline and enhancing tissue regeneration in aging populations.

## Materials and Methods

### Experimental design

The primary objective of this research was to characterize age-related changes within the human non-hematopoietic BM microenvironment using single-cell transcriptomics. Our initial hypothesis was that significant transcriptional alterations within BM stromal cell populations drive BM aging. To test this, we used human BM samples obtained from healthy young and elderly individuals undergoing orthopedic surgery. Key discoveries arising from the analysis of the transcriptomic data were subsequently validated through IF and spatial transcriptomics. The specific experimental workflow, including detailed information on sample origin and processing, as well as the statistical methods employed, is described in the subsequent sections.

### Human samples

Human BM samples were obtained from BM tissue of healthy young (28-48 years of age) and elderly (58-79 years of age) individuals of both sexes undergoing orthopedic surgery (hip or knee replacement). Samples were collected at Hospital Universitario de Navarra and Hospital Reina Sofía de Tudela. Written informed consent was obtained according to the Declaration of Helsinki and the Institutional Review Board of the University of Navarra. The characteristics of all individuals are described in **Table S1**.

### Isolation and fluorescence-activated cell sorting of endothelial and mesenchymal BM cells

Endothelial and mesenchymal human non-hematopoietic BM cells were isolated and sorted for subsequent scRNA-seq analysis (**Fig S1**). All sample processing steps were performed on ice to preserve cell viability and RNA integrity, except red blood cell lysis, which was conducted at room temperature (RT). To isolate endothelial and mesenchymal BM cells, red blood cells were lysed using a 45 ml:5 ml ratio of ACK lysis buffer per human sample, filtered through a 70 μm filter, and then collected into a tube. Bone fragments were manually cut, crushed, and digested with 0.3% collagenase I and dispase (5U/ml) for 15 min at 37°C and shaken at 200 rpm. After digestion, the collagenized bone fraction was filtered through a 70 μm filter into a collection tube, pooled with BM fraction, and centrifuged at 1500rpm and 4°C for 5 minutes. To determine live cell concentration and viability, 10 μl of each sample was stained with acridine orange and propidium iodide (AO/PI) solution (Nexcelom) and analyzed with Cellometer K2 Image Cytometer (Nexcelom Bioscience). Cells were subsequently stained for 30min on ice in PBS 1X containing 2 % FBS and 2 mM EDTA (modPBS) with the following combination of conjugated antibodies: BV510 labelled anti-Lin (including CD3, CD19, CD45 and CD64), BV421 labelled anti-CD235, BV421 labelled anti-CD45, FITC labelled anti-CD16, PE-Cy7 labelled anti-CD56, PE labelled anti-CD31, APC-Cy7 labelled anti-CD9, and PerCP-Cy5.5 labelled anti-CD271. All antibodies were added at a concentration of 1/100 except anti-Lin, where we added 3 μl/test-test 25×106cells, anti-CD16, 10 μl/test-test 25×106cells and 1/50 anti-CD31 and anti-CD56. Samples were resuspended in modPBS with 6 μl of TO-PRO-3 for cell sorting. Sorting gates were set according to the corresponding fluorescence-minus-one (FMO) controls, and cells were sorted using BD FACSAria II. Dead cells were excluded by TO-PRO-3 staining, and doublets and debris were excluded by gating on FSC and SSC. EC and MSC were prospectively isolated based on the following immunophenotype (**Fig S1**): TO-PRO-3-/Lin-/CD45-/CD235-/CD16-/CD56-/CD9+/CD31+ for EC and TO-PRO-3-/Lin-/CD45-/CD235-/CD16-/CD56-/CD31-/CD271+ for MSC. EC and MSC were directly sorted into PBS 1x with 0.05% UltraPure for subsequent scRNA-seq protocol. Cell viability of sorted cells was assessed using Nexcelom Cellometer as described above.

### Single-cell RNA sequencing

scRNA-seq was performed using the Single Cell 3’ Reagent Kits v3.1 (10X Genomics) according to the manufacturer’s instructions. Approximately 10,000 cells were loaded at a concentration of 1,000 cells/µL on a Chromium Controller instrument (10X Genomics) to generate single-cell gel bead-in-emulsions (GEMs). Each cell was encapsulated with primers containing a fixed Illumina Read one sequence, followed by a cell-identifying 16 bp 10X barcode, a 10 bp Unique Molecular Identifier (UMI), and a poly-dT sequence. A subsequent reverse transcription yielded full-length, barcoded cDNA. This cDNA was then released from the GEMs, PCR-amplified, and purified with magnetic beads (SPRIselect, Beckman Coulter). Enzymatic Fragmentation and Size Selection optimized cDNA size before library construction. Illumina adaptor sequences were added, and the resulting library was amplified via end repair, A-tailing, adaptor ligation, and PCR. Libraries’ quality control and quantification were performed using Qubit 3.0 Fluorometer (Life Technologies) and Agilent’s 4200 TapeStation System (Agilent), respectively. Sequencing was performed in a NextSeq2000 (Illumina) (Read 1: 28cycles, i7 Index: 10cycles, i5 Index:10 Read 2: 90cycles) at an average depth of 45,000 reads/cell followed by computational alignment using CellRanger (version (v) 6.1.1, 10x Genomics) against the human GRCh38 reference genome.

### Single-cell RNA sequencing analysis

The single-cell analysis of the BM samples was performed using R version (v) 4.1.3 and Seurat v 4.3.0(*73*). For data pre-processing, doublet scores were calculated by scDblFinder v.1.8.0(*74*). Datasets were filtered individually based on a library complexity of more than 200 features, features detected in more than 3 cells, doublets, and high percentages of mitochondrial genes (>10%). After individual sample pre-processing, single-cell datasets from young and old individuals were merged, normalized using the SCTransform method(*75*), and analyzed by principal-component analysis (PCA) (k = 30) on the most variable genes (k = 3,000) across all cells. A batch correction was performed using the RunHarmony function from the Harmony R package(*76*), setting the sample source as the batch factor. The batch-corrected coordinate space was used for linear dimensional reduction with Seurat. Unsupervised clustering was performed by computing the K-nearest neighbors, applying the Louvain algorithm at resolution 0.4, and cells were projected in two dimensions using Uniform Manifold Approximation and Projection (UMAP).

Significantly upregulated genes in each cluster compared to all other clusters (Bonferroni-adjusted p-values <0.05) were identified using the Seurat Function FindMarkers following MAST (Model-based Analysis of Single-cell Transcriptomics) methodology(*77*). Cell types were identified by manual cell type annotations according to published canonical marker genes.

### Characterization of sub-cell types and functional states in human BM EC and MSC

To characterize endothelial and mesenchymal BM compartments, clusters identified as EC and MSC were subsetted into separate objects, and unsupervised cell clustering was implemented. The stability of the clusters was evaluated following a bootstrapping strategy, and cells from non-robust clusters were assigned to the neighboring clusters, repeating a Random-Forest-based strategy as described in our previous study(*18*). Additionally, two clusters identified within EC and some outlier cells within MSC, lacking canonical marker genes for their respective population, were excluded from downstream analysis. To identify the markers for each cluster, genes that were differentially expressed (min. pct > 0.25, log fold change > 1; and Bonferroni adjusted p-value < 0.05) in a cluster compared to all the others were identified using the FindAllMarkers function with MAST. GO ORA was conducted using clusterProfiler to detect enriched canonical pathways (adjusted p-value < 0.05) among the identified cluster markers (*26*, *78*). Cell clustering identities were then annotated by cross-referencing these cluster-specific genes and functional gene sets with published data.

### Age-related differential expression and functional enrichment analysis

DEGs between young and aged cells were determined with the Seurat FindMarkers function with MAST (min. pct > 0.25, absolute log fold change > 0.5; and Bonferroni adjusted p-value < 0.05). Significant DEGs for each group were used as input for a GO ORA using clusterProfiler to investigate the biological differences between age groups. GO terms with corrected P-values less than 0.05 were considered significantly enriched. GSEA was also calculated using the GO and Hallmark gene sets from the Molecular Signatures Database v.7.5.1 in the list of DEGs ranked according to log2FC.

### Age-related changes in cell proportion

To robustly determine the statistical significance of differences in cell-type proportions between young and aged conditions we conducted a permutation-based analysis using the R-package scProportionTest(*79*). Briefly, this method runs a permutation test on the two conditions for each cell cluster. It returns the relative change in cell type abundance between the two groups with a confidence interval for each comparison. To account for inter-individual variability in cellular composition and to improve the robustness of confidence interval estimation, we additionally applied a bootstrapping strategy. In each iteration, an equal number of cells from each age group was randomly selected. For each cell type in each iteration, the number of cells was determined and divided by the total cell number of the iteration to calculate the cell-type proportions. Bootstrap sampling was used to generate 95% confidence intervals for plotting. Changes in population sizes with FDR < 0.05 and log2FC > 1 were denoted as statistically significant.

### Gene Regulatory Network analysis

GRN activity was interrogated following the workflow implemented by SCENIC(*30*). An equal number of cells per condition and cell type was randomly selected for the analysis. Briefly, all the genes were trained in the GENIE3 package and used to develop a random forest model for selecting co-expression modules between TFs and target genes (regulons). Regions for TFs searching were restricted to a 10-k distance centered on the transcriptional start site (TSS) or 500 bp upstream of the TSSs. Then, RcisTarget was used to refine the regulons by inferring direct targets of the TFs. Finally, regulon activity scores were calculated for each cell, using the AUCell package, to determine whether the regulons were in an active or inactive state.

SCENIC GRN results were further validated using SimiC(*31*). For each comparison, 100 TFs and 1,000 target genes were selected based on their variability, determined by calculating the maximum absolute deviation. To determine the optimal parameters, each analysis involved a cross-validation run. After parameter tuning with cross-validation, we set lambda1 = 0.01, lambda2 = 0.1. We extracted regulons by calculating association weights for each TF and target and filtering out small weights. The resulting GRNs were visualized using the GRN incidence matrices provided by SimiC. Histograms for different regulons were computed using the “regulon activity score” provided by SimiC. We then computed the weighted area under the curve (wAUC) to measure regulon activity per cell. Additionally, this score was utilized to calculate the regulatory dissimilarity score between functional cell states. Lastly, we tested for differences in regulon activity between young and old EC and MSC, performing a Kolmogorov–Smirnov test for the wAUC distributions.

### Cell-to-cell communication analysis

An equal number of cells per condition and cell type was randomly selected for the analysis to predict cell-cell interactions between the endothelial, mesenchymal, and hematopoietic populations from the BM microenvironment. The liana R package v.0.1.13(*44*) was used to predict cell-cell interactions using five methods (cellphonedb, connectome, NATMI, logFC, and SingleCellSignalR). The liana_aggregate function was used to generate an aggregate rank score for each interaction, reflecting only the specificity of interactions, and interactions were filtered based on their statistical significance (p-value < 0.01). To illustrate the strength of specific interactions between different groups, several L-R pairs with high mean values were selected for visualization. The complete list of ligand-receptor interaction pairs is shown in **Table S12.**

### Human BM biopsies

Human BM biopsies from young (34-48 years of age) and aged (72-78 years of age) individuals were obtained from Hospital Universitario de Navarra after obtaining written informed consent for IF and spatial analysis (**Table S9)**. Archived paraffined human BM biopsies of four MF patients (65–73 years of age) were obtained from Pathology Department of Clínica Universidad de Navarra after written informed consent was achieved (**Table S9)**. The Research Ethics Committee of the University of Navarra approved the human sample collection and research conducted in this study. Personal data was kept confidential following the Organic Law 3/2018 on personal data protection and Spanish Law 14/2007 on Biomedical research. All collection samples are codified; only authorized personnel can correlate the patient’s identity with the codes.

### Immunofluorescence and Masson’s Trichrome staining

Bone tissue sections were fixed in formol 4% (PanReac) for 24 hours at RT and then decalcified using EDTA 0.25 M pH 6.95 (Invitrogen) for 10 days at RT with agitation. Following decalcification, the samples were washed in distilled water for 5 min and sequentially dehydrated in ethanol at increasing concentrations: 70% for 1 hour, 80% for 1 hour, 96% for 1 hour, and 100% overnight. The samples were then cleared in xylol (PanReac) for 4 hours. Fixed bone samples were subsequently embedded in paraffin and incubated at 60 °C overnight. Subsequently, 4 µm tissue sections were mounted on microscopy slides and dried in a desiccator at 37 °C overnight. The preparations were deparaffinized and rehydrated through a series of graded alcohols. Antigen retrieval was performed by heating the slides in 10 mM Tris-1 mM EDTA buffer (pH 9) for 30 minutes at 95°C, except samples for RAB13 and EMCN staining that were heated in 10 µM Citrate (pH6).

For Masson’s Trichrome staining, the samples were performed according to standard protocols to determine fibrosis in the human BM niche(*80*). Tissue imaging was performed using an Aperio CS2 Scanner (Leica Biosystems) at 20x magnification. Image analysis was achieved using QuPath (version 0.5.1) to evaluate cell circularity and ImageJ (version 2.14.0) to quantify fiber percentage based on color intensity.

For IF staining, the tissue sections were blocked with BSA 5% for 30 min at RT and the following primary antibodies were used: rabbit anti-CD45 Alexa Fluor 647 conjugate (Cell Signaling Technology, 1:50 dilution), mouse anti-CD31 antibody (DakoCytomation, 1:40 dilution), rabbit anti-PRRX1 antibody (Sigma-Aldrich, 1:50 dilution), mouse anti-Alpha smooth muscle actin antibody (Sigma Aldrich 1:1000 dilution), rat anti-Endomucinantibody (V.7C7) (Santa Cruz Technology 1: 40 dilution), mouse PE anti-human CD144 (VE-Cadherin) Antibody (BioLegend 1:50 dilution), mouse Anti-human RAB13 Antibody (MA5-31879) (ThermoFisher Scientific 1:100 dilution), mouse anti-human BGLAP antibody (Sigma Aldrich 1:100 dilution) and mouse anti-human CD90 Antibody (MA5-16671) (ThermoFisher Scientific 1:100 dilution). The slides were subsequently incubated with the corresponding secondary antibodies in a 1:200 dilution: goat anti-Rabbit IgG Alexa Fluor™ 647 (Invitrogen), goat anti-Mouse Alexa Fluor™ 568 (Invitrogen), goat anti-Mouse Alexa Fluor™ 488 (Invitrogen), and goat anti-Rat Alexa Fluor™ 647 (Invitrogen). Nuclei were stained with DAPI (Vectashield 1:50 dilution). Fluorescence images were acquired using the Vectra Polaris Multispectral Imaging System (Perkin Elmer). Image analysis was made using QuPath (0.5.1) to assess the co-expression of CD90 and PRRX1 in the BM of young, elderly, and MF patients.

### Preparation and Sequencing of Spatial Transcriptomics Samples and Libraries

Human bone samples were processed as previously described(*71*). Bone tissue sections were fixed in formol 4% (PanReac) for 24 h at RT and then decalcified with EDTA 0.25 M pH 6.95 (Invitrogen) for 10 days at RT with agitation. After decalcification, samples were washed in distilled water for 5 min and sequentially dehydrated in ethanol 70% (1 h), 80% (1 h), 96% (1 h), and 100% (overnight), followed by 4 h in xylol (Panreac). Fixed bone samples were subsequently embedded in paraffin and incubated at 60 °C overnight.

Prior to spatial transcriptomics experiments, RNA integrity of the FFPE tissue, a critical factor for spatial transcriptomics success, was examined on 10 µm sections. RNA was extracted using RNeasy FFPE kit (Qiagen) and examined with RNA ScreenTape Assay to determine the samples DV200. Next, 5 μm tissue sections were mounted on microscopy slides, baked at 42 °C for 3 h, and dried in a desiccator at 37 °C overnight. For spatial transcriptomics assays, preparations were deparaffined, rehydrated, and H&E stained following 10X Genomics recommendations. Tissue imaging was performed using an Aperio CS2 Scanner (Leica Biosystems) at 20x magnification. Sections were destained and decrosslinked before library construction using Visium CytAssist Spatial Gene Expression for FFPE Human Transcriptome (10X Genomics). Briefly, a whole transcriptome panel consisting of 3 probe pairs per gene was hybridized with their target RNAs in the tissue sections. Neighboring probe pairs that had hybridized to RNA were then ligated. Tissue slides and Visium CytAssist Spatial Gene Expression v2 Slides were loaded into a Visium CytAssist instrument (10X Genomics). Through RNase treatment and tissue permeabilization, ligated probes were released and readily diffused onto Visium v2 slides containing spatially barcoded oligonucleotides. Probes were spatially labeled through extension, released from the slide, and pooled. Finally, samples were indexed via PCR amplification. The resulting libraries were quantified with Qubit dsDNA HS Assay Kit, and their profile was examined using Agilent’s HS D1000 ScreenTape Assay. Sequencing was carried out in an Illumina NextSeq2000 using paired-end, dual-index sequencing (Rd1: 28 cycles; i7: 10 cycles; i5: 10 cycles; Rd2: 50 cycles) at a minimum depth of 25000 reads per spot.

### Pre-processing and QC spatial transcriptomics data

Spatial transcriptomic data were demultiplexed and mapped using the SpaceRanger software (v2.0.1). The Ensembl 105 genomes were used as reference (GRCh38). Filtered feature-barcode expression matrices obtained from SpaceRanger were used as initial input for the spatial transcriptomics analysis using Seurat (v4.3.0.1) and Stutility (v1.1.1). Spots overlapping the bone tissue were manually removed based on the H&E staining images using the Cloupe (v6.0.0). Spots with less than 200 UMI and 150 features were filtered out. Immunoglobulin genes and those genes that were not present in both samples were filtered out.

### Deconvolution analysis

We downloaded a pre-processed human single-cell dataset published by Bandyopadhyay et al.(*21*) for deconvolution analysis. We removed or relabeled their cell type annotation to obtain our interest’s 10 major cell types (B-cells, T-cells, Neutrophils, Erythroblasts, Monocytes, DC, PC, MSC, EC, and HSC). Deconvolution analysis used the relabeled single-cell reference with CARD (v1.0.0) with default parameters.

### Spot labeling

The estimated cell-type percentage values derived from the deconvolution were used to define the percentage of each cell type in each sample. We used the deconvolution estimation and the area under the curve (AUC) calculated for each cell type per spot to label the spots. We first obtained the top marker genes for each cell type by comparing the transcriptome versus the rest of the cell types included in our reference using the default Seurat pipeline. Cell type AUC scores were calculated using AUCell (v1.18.1), which computes the area under the curve for a given gene set (a unitless measure) per spot. This score, also called signature, reflects the activity of a gene set. For each cell type, we ranked the spots based on the AUC score and the estimated percentage value. Following these criteria, each spot was labeled as golden, high, low, or rest for the specific cell type.

- First, we defined the average percentage of each cell type based on the deconvolution analysis: e.g. 32.2% for MSC and 16.0% for EC.
- Spots among the top 32.2% for MSC (16.0% for EC) in both AUC and deconvolution rankings were labeled as Golden.
- Spots among the top 64.4% for MSC (32.0% for EC) in both rankings, which are not Golden, were labeled High.
- Spots with an AUC = 0, independently of the deconvolution percentage, were labeled as rest.
- The remaining spots were labeled as <u>Low</u>.

MSC and EC golden spots were then used for deconvolution and AUC analysis of the specific subtypes using the single-cell data generated in the study.

### Hierarchical clustering

The estimated cell-type percentage values derived from the deconvolution were further used for hierarchical clustering with ISCHIA. Samples were merged, and the optimal number of clusters was 7. The cell-type prevalence within each cluster was then calculated. The prevalence of each cluster in each sample was determined.

### Cell type correlation

To determine whether two cell types co-localize within the same spots, we conducted a correlation analysis of their AUC scores across the spots of interest (“golden” spots for the cell type of study). A Spearman correlation test was performed to assess the significance of this correlation. Additionally, to evaluate whether the correlation between the two cell types changes with distance from the spots of interest, we calculated the average AUC score in the six nearest spots surrounding each spot of interest. The AUC score of the target cell type within its respective “golden” spot was then correlated with the average AUC score of the other cell types in the surrounding spots. A Spearman correlation test was again used to assess the significance of this correlation.

### Statistical analyses

Statistical analysis was carried out in R (v4.1.3). Tests used to evaluate statistical significance are detailed in each method section.

## Acknowledgments

We would like to thank the staff of the flow cytometry core and the advanced genomic lab for their invaluable technical and intellectual assistance. We also acknowledge Hospital Universitario de Navarra and Hospital Reina Sofía de Tudela for providing the human BM samples and their collaboration. We are particularly grateful to the healthy donors for participating in this study.

## Funding

Instituto de Salud Carlos III and co-financed by ERDF A way of making Europe (PI20/01308, PI23/00516)

CIBERONC (CB16/12/00489), RICORS TERAV and TERAV Plus (RD21/0017/0009) (RD24/0014/0010)

Departamento de Industria Gobierno de Navarra (AGATA 0011-1411-2020-000010/0011-1411-2020-000011)

Departamento de Salud Gobierno de Navarra to F.P Spanish Government through project PID2019-111192GA-I00 (MICINN) Marie Curie grant from the European Commission (H2020-MSCA-IF-837491) AECC predoctoral fellowship (PRDNA19006CENZ)

FPU fellowship from Ministerio de Ciencia, Innovación y Universidades (FPU22/03283)

## Author contributions

I.A.C., I.C. and M.C. processed human BM samples. I.C. analyzed the scRNA-seq data and performed the cell-to-cell interactome. A.R.L.P. analyzed the spatial transcriptomic data. J.R., I.S.G., M.M.B., and D.Q.A. provided the BM samples. P.S.M., P.A.R., and A.V. performed the scRNA-seq experiments. D.A. and A.L. performed the fluorescence-activated cell sorting. I.A.C., I.C., and M.C. interpreted the data and wrote the paper. S.S., P.R.C., P.S.M., and P.A.R. performed the spatial transcriptomic experiments. M.C., and L.C-D. performed the histologic analysis. D.G.C., with help from J.Y., L.S., R.L., and J.T., supervised the computational work. M.A-M. and E.M. contributed by providing the myelofibrosis samples. I.A.C. and B.S. conceptualized the study. F.P., D.G.C., and I.A.C conceived and directed the research project. All authors actively participated in the discussions underlying this manuscript. F.P., D.G.C., I.A.C, I.C., and M.C. discussed the results and wrote the final manuscript. All authors contributed to, read, and approved the final manuscript.

## Competing interests

### Data and materials availability

The scRNA-seq data generated in this study and spatial data from the young sample will be publicly available as of the date of peer-reviewed publication and is available under request, contact. Spatial raw and processed data from the elderly sample has been deposited at GSE269875. scRNA-seq data used for spatial deconvolution was obtained from GSE253355.

## References

1. C. J. Eaves, Hematopoietic stem cells: Concepts, definitions, and the new reality. Blood 125, 2605–2613 (2015).

2. R. Schofield, The relationship between the spleen colony-forming cell and the haemopoietic stem cell. Blood Cells 4, 7–25 (1978).

3. S. J. Morrison, D. T. Scadden, The bone marrow niche for haematopoietic stem cells. Nature 505, 327–334 (2014).

4. S. Pinho, P. S. Frenette, Haematopoietic stem cell activity and interactions with the niche. Nat Rev Mol Cell Biol 20, 303–320 (2019).

5. J. Fröbel, T. Landspersky, G. Percin, C. Schreck, S. Rahmig, A. Ori, D. Nowak, M. Essers, C. Waskow, R. A. J. Oostendorp, The Hematopoietic Bone Marrow Niche Ecosystem. Front Cell Dev Biol 9 (2021).

6. S. S. Khan, B. D. Singer, D. E. Vaughan, Molecular and physiological manifestations and measurement of aging in humans. Aging Cell 16, 624–633 (2017).

7. T. Y. Su, J. Hauenstein, E. Somuncular, Ö. Dumral, E. Leonard, C. Gustafsson, E. Tzortzis, A. Forlani, A. S. Johansson, H. Qian, R. Månsson, S. Luc, Aging is associated with functional and molecular changes in distinct hematopoietic stem cell subsets. Nature Communications 15 (2024).

8. J. Guo, X. Huang, L. Dou, M. Yan, T. Shen, W. Tang, J. Li, Aging and aging-related diseases: from molecular mechanisms to interventions and treatments. Signal Transduct Target Ther 7, 391 (2022).

9. P. Ramalingam, M. C. Gutkin, M. G. Poulos, T. Tillery, C. Doughty, A. Winiarski, A. G. Freire, S. Rafii, D. Redmond, J. M. Butler, Restoring bone marrow niche function rejuvenates aged hematopoietic stem cells by reactivating the DNA Damage Response. Nature Communications 2023 14:*1* **14**, 1–20 (2023).

10. M. G. Poulos, P. Ramalingam, M. C. Gutkin, P. Llanos, K. Gilleran, S. Y. Rabbany, J. M. Butler, Endothelial transplantation rejuvenates aged hematopoietic stem cell function. Journal of Clinical Investigation 127, 4163–4178 (2017).

11. Y.-H. Ho, S. Méndez-Ferrer, Microenvironmental contributions to hematopoietic stem cell aging. Haematologica 105, 38–46 (2020).

12. S. Dobner, F. Tóth, L. P. M. H. de Rooij, A high-resolution view of the heterogeneous aging endothelium. Angiogenesis 2024 27:*2* **27**, 129–145 (2024).

13. M. Al-Azab, M. Safi, E. Idiiatullina, F. Al-Shaebi, M. Y. Zaky, Aging of mesenchymal stem cell: machinery, markers, and strategies of fighting. Cellular & Molecular Biology Letters 2022 27:*1* **27**, 1–40 (2022).

14. T. Fujino, S. Asada, S. Goyama, T. Kitamura, Mechanisms involved in hematopoietic stem cell aging. Cellular and Molecular Life Sciences 79(9):473 (2022). 10.1007/s00018-022-04356-5.

15. P. M. Helbling, E. Piñeiro-Yáñez, R. Gerosa, S. Boettcher, F. Al-Shahrour, M. G. Manz, C. Nombela-Arrieta, Global Transcriptomic Profiling of the Bone Marrow Stromal Microenvironment during Postnatal Development, Aging, and Inflammation. Cell Rep 29, 3313–3330.e4 (2019).

16. W. Chen, X. Chen, L. Yao, J. Feng, F. Li, Y. Shan, L. Ren, C. Zhuo, M. Feng, S. Zhong, C. He, A global view of altered ligand-receptor interactions in bone marrow aging based on single-cell sequencing. Comput Struct Biotechnol J 23, 2754–2762 (2024).

17. Z. Wang, X. Li, J. Yang, Y. Gong, H. Zhang, X. Qiu, Y. Liu, C. Zhou, Y. Chen, J. Greenbaum, L. Cheng, Y. Hu, J. Xie, X. Yang, Y. Li, M. R. Schiller, Y. Chen, L. Tan, S.-Y. Tang, H. Shen, H.-M. Xiao, H.-W. Deng, Single-cell RNA sequencing deconvolutes the in vivo heterogeneity of human bone marrow-derived mesenchymal stem cells. *Ivyspring International Publisher Int*. J. Biol. Sci 2021, 4192–4206 (2021).

18. J. Ye, I. A. Calvo, I. Cenzano, A. Vilas, X. Martinez-de-Morentin, M. Lasaga, D. Alignani, B. Paiva, A. C. Viñado, P. San Martin-Uriz, J. P. Romero, D. Quilez Agreda, M. Miñana Barrios, I. Sancho-González, G. Todisco, L. Malcovati, N. Planell, B. Saez, J. N. Tegner, F. Prosper, D. Gomez-Cabrero, Deconvolution of the hematopoietic stem cell microenvironment reveals a high degree of specialization and conservation. iScience 25 (2022).

19. H. Li, S. Bräunig, P. Dhapolar, G. Karlsson, S. Lang, S. Scheding, Identification of phenotypically, functionally, and anatomically distinct stromal niche populations in human bone marrow based on single-cell RNA sequencing. Elife 12 (2023).

20. L. Fiévet, N. Espagnolle, D. Gerovska, D. Bernard, C. Syrykh, C. Laurent, P. Layrolle, J. De Lima, A. Justo, N. Reina, L. Casteilla, M. J. Araúzo-Bravo, A. Naji, J. C. Pagès, F. Deschaseaux, Single-cell RNA sequencing of human non-hematopoietic bone marrow cells reveals a unique set of inter-species conserved biomarkers for native mesenchymal stromal cells. Stem Cell Res Ther 14, 1–16 (2023).

21. S. Bandyopadhyay, M. P. Duffy, K. J. Ahn, J. H. Sussman, M. Pang, D. Smith, G. Duncan, I. Zhang, J. Huang, Y. Lin, B. Xiong, T. Imtiaz, C. H. Chen, A. Thadi, C. Chen, J. Xu, M. Reichart, Z. Martinez, C. Diorio, C. Chen, V. Pillai, O. Snaith, D. Oldridge, S. Bhattacharyya, I. Maillard, M. Carroll, C. Nelson, L. Qin, K. Tan, Mapping the cellular biogeography of human bone marrow niches using single-cell transcriptomics and proteomic imaging. Cell 187, 3120–3140.e29 (2024).

22. I. L. Boueya, L. Sandhow, J. R. P. Albuquerque, R. Znaidi, D. Passaro, Endothelial heterogeneity in bone marrow: insights across development, adult life and leukemia. Leukemia 2024, 1–17 (2024).

23. J. Kalucka, L. P. M. H. de Rooij, J. Goveia, K. Rohlenova, S. J. Dumas, E. Meta, N. V. Conchinha, F. Taverna, L. A. Teuwen, K. Veys, M. García-Caballero, S. Khan, V. Geldhof, L. Sokol, R. Chen, L. Treps, M. Borri, P. de Zeeuw, C. Dubois, T. K. Karakach, K. D. Falkenberg, M. Parys, X. Yin, S. Vinckier, Y. Du, R. A. Fenton, L. Schoonjans, M. Dewerchin, G. Eelen, B. Thienpont, L. Lin, L. Bolund, X. Li, Y. Luo, P. Carmeliet, Single-cell transcriptome atlas of murine endothelial cells. Cell 180, 764–779.e20 (2020).

24. S. N. Barnett, A.-M. Cujba, L. Yang, A. R. Maceiras, S. Li, V. R. Kedlian, J. P. Pett, K. Polanski, A. M. A. Miranda, C. Xu, J. Cranley, K. Kanemaru, M. Lee, L. Mach, S. Perera, C. Tudor, P. D. Joseph, S. Pritchard, R. Toscano-Rivalta, Z. K. Tuong, L. Bolt, R. Petryszak, M. Prete, B. Cakir, A. Huseynov, I. Sarropoulos, R. A. Chowdhury, R. Elmentaite, E. Madissoon, A. J. Oliver, L. Campos, A. Brazovskaja, T. Gomes, B. Treutlein, C. N. Kim, T. J. Nowakowski, K. B. Meyer, A. M. Randi, M. Noseda, S. A. Teichmann, An organotypic atlas of human vascular cells. Nat Med 30, 3468 (2024).

25. A. Thiriot, C. Perdomo, G. Cheng, I. Novitzky-Basso, S. McArdle, J. K. Kishimoto, O. Barreiro, I. Mazo, R. Triboulet, K. Ley, A. Rot, U. H. von Andrian, Differential DARC/ACKR1 expression distinguishes venular from non-venular endothelial cells in murine tissues. BMC Biol 15, 1–19 (2017).

26. M. Ashburner, C. A. Ball, J. A. Blake, D. Botstein, H. Butler, J. M. Cherry, A. P. Davis, K. Dolinski, S. S. Dwight, J. T. Eppig, M. A. Harris, D. P. Hill, L. Issel-Tarver, A. Kasarskis, S. Lewis, J. C. Matese, J. E. Richardson, M. Ringwald, G. M. Rubin, G. Sherlock, Gene Ontology: tool for the unification of biology. Nature Genetics 2000 25:*1* **25**, 25–29 (2000).

27. S. Hopkin, J. M. Lord, M. Chimen, Dysregulation of leukocyte trafficking in ageing: Causal factors and possible corrective therapies. Pharmacol Res 163, 105323 (2021).

28. N. Wechter, M. Rossi, C. Anerillas, D. Tsitsipatis, Y. Piao, J. Fan, J. L. Martindale, S. De, K. Mazan-Mamczarz, M. Gorospe, Single-cell transcriptomic analysis uncovers diverse and dynamic senescent cell populations. Aging (Albany NY) 15, 2824 (2023).

29. A. Varesi, L. I. M. Campagnoli, A. Barbieri, L. Rossi, G. Ricevuti, C. Esposito, S. Chirumbolo, N. Marchesi, A. Pascale, RNA binding proteins in senescence: A potential common linker for age-related diseases? Ageing Res Rev 88 (2023).

30. S. Aibar, C. B. González-Blas, T. Moerman, V. A. Huynh-Thu, H. Imrichova, G. Hulselmans, F. Rambow, J. C. Marine, P. Geurts, J. Aerts, J. Van Den Oord, Z. K. Atak, J. Wouters, S. Aerts, SCENIC: Single-cell regulatory network inference and clustering. Nat Methods 14, 1083 (2017).

31. J. Peng, G. Serrano, I. M. Traniello, M. E. Calleja-Cervantes, U. V. Chembazhi, S. Bangru, T. Ezponda, J. R. Rodriguez-Madoz, A. Kalsotra, F. Prosper, I. Ochoa, M. Hernaez, SimiC enables the inference of complex gene regulatory dynamics across cell phenotypes. Communications Biology 2022 5:*1* **5**, 1–19 (2022).

32. A. Almotiri, A. Abdelfattah, E. Storch, M. P. Stemmler, S. Brabletz, T. Brabletz, N. P. Rodrigues, Zeb1 maintains long-term adult hematopoietic stem cell function and extramedullary hematopoiesis. Exp Hematol 134 (2024).

33. Q. C. Yu, A. Geng, C. B. Preusch, Y. Chen, G. Peng, Y. Xu, Y. Jia, Y. Miao, H. Xue, D. Gao, L. Bao, W. Pan, J. Chen, K. C. Garcia, T. H. Cheung, Y. A. Zeng, Activation of Wnt/β-catenin signaling by Zeb1 in endothelial progenitors induces vascular quiescence entry. Cell Rep 41 (2022).

34. D. Akrivou, G. Perlepe, P. Kirgou, K. I. Gourgoulianis, F. Malli, Pathophysiological Aspects of Aging in Venous Thromboembolism: An Update. Medicina (Kaunas) 58 (2022).

35. M. B. Chen, A. C. Yang, B. Lehallier, S. R. Quake, T. Wyss-Coray, Brain Endothelial Cells Are Exquisite Sensors of Age-Related Circulatory Cues. doi: 10.1016/j.celrep.2020.03.012 (2020).

36. M. J. Zhang, A. O. Pisco, S. Darmanis, J. Zou, Mouse aging cell atlas analysis reveals global and cell type-specific aging signatures. Elife 10 (2021).

37. S. Stucker, J. Chen, F. E. Watt, A. P. Kusumbe, Bone Angiogenesis and Vascular Niche Remodeling in Stress, Aging, and Diseases. Front Cell Dev Biol 8, 602269 (2020).

38. Y. Tian, Y. Tian, Z. Yuan, Y. Zeng, S. Wang, X. Fan, D. Yang, M. Yang, Iron Metabolism in Aging and Age-Related Diseases. Int J Mol Sci 23, 3612 (2022).

39. K. Ghosh, D. K. Shome, B. Kulkarni, M. K. Ghosh, K. Ghosh, Fibrosis and bone marrow: understanding causation and pathobiology. J Transl Med 21, 703 (2023).

40. L. Longhitano, D. Tibullo, N. Vicario, C. Giallongo, E. La Spina, A. Romano, S. Lombardo, M. Moretti, F. Masia, A. R. D. Coda, S. Venuto, P. Fontana, R. Parenti, G. L. Volti, M. Di Rosa, G. A. Palumbo, A. Liso, IGFBP-6/sonic hedgehog/TLR4 signalling axis drives bone marrow fibrotic transformation in primary myelofibrosis. Aging (Albany NY) 13, 25055 (2021).

41. D. Duscher, R. C. Rennert, M. Januszyk, E. Anghel, Z. N. Maan, A. J. Whittam, M. G. Perez, R. Kosaraju, M. S. Hu, G. G. Walmsley, D. Atashroo, S. Khong, A. J. Butte, G. C. Gurtner, Aging disrupts cell subpopulation dynamics and diminishes the function of mesenchymal stem cells. doi: 10.1038/srep07144.

42. Y. Zhang, L. Guo, S. Han, L. Chen, C. Li, Z. Zhang, Y. Hong, X. Zhang, X. Zhou, D. Jiang, X. Liang, J. Qiu, J. Zhang, X. Li, S. Zhong, C. Liao, B. Yan, H.-F. Tse, Q. Lian, Adult mesenchymal stem cell ageing interplays with depressed mitochondrial Ndufs6. Cell Death Dis 11, 1075 (2020).

43. D. Ghosh, C. Mejia-Pena, N. Quach, B. Xuan, A. H. Lee, M. R. Dawson, Senescent mesenchymal stem cells remodel extracellular matrix driving breast cancer cells to more invasive phenotype. J Cell Sci, doi: 10.1242/jcs.232470 (2020).

44. D. Dimitrov, D. Türei, M. Garrido-Rodriguez, P. L. Burmedi, J. S. Nagai, C. Boys, R. O. Ramirez Flores, H. Kim, B. Szalai, I. G. Costa, A. Valdeolivas, A. Dugourd, J. Saez-Rodriguez, Comparison of methods and resources for cell-cell communication inference from single-cell RNA-Seq data. Nature Communications 2022 13:*1* **13**, 1–13 (2022).

45. A. Akil, A. K. Gutiérrez-García, R. Guenter, J. B. Rose, A. W. Beck, H. Chen, B. Ren, Notch Signaling in Vascular Endothelial Cells, Angiogenesis, and Tumor Progression: An Update and Prospective. Front Cell Dev Biol 9, 642352 (2021).

46. K. R. Brunt, Y. Zhang, A. Mihic, M. Li, S. H. Li, P. Xue, W. Zhang, S. Basmaji, K. Tsang, R. D. Weisel, T. M. Yau, R. K. Li, Role of WNT/β-Catenin Signaling in Rejuvenating Myogenic Differentiation of Aged Mesenchymal Stem Cells from Cardiac Patients. Am J Pathol 181, 2067–2078 (2012).

47. V. Donega, A. T. van der Geest, J. A. Sluijs, R. E. van Dijk, C. C. Wang, O. Basak, R. J. Pasterkamp, E. M. Hol, Single-cell profiling of human subventricular zone progenitors identifies SFRP1 as a target to re-activate progenitors. Nature Communications 2022 13:*1* **13**, 1–12 (2022).

48. A. Agrawal, S. Agrawal, S. Gupta, Role of Dendritic Cells in Inflammation and Loss of Tolerance in the Elderly. Front Immunol 8 (2017).

49. P. L. Ståhl, F. Salmén, S. Vickovic, A. Lundmark, J. F. Navarro, J. Magnusson, S. Giacomello, M. Asp, J. O. Westholm, M. Huss, A. Mollbrink, S. Linnarsson, S. Codeluppi, Å. Borg, F. Pontén, P. I. Costea, P. Sahlén, J. Mulder, O. Bergmann, J. Lundeberg, J. Frisén, Visualization and analysis of gene expression in tissue sections by spatial transcriptomics. Science (1979) 353, 78–82 (2016).

50. M. Ainciburu, T. Ezponda, N. Berastegui, A. Alfonso-Pierola, A. Vilas-Zornoza, P. S. Martin-Uriz, D. Alignani, J. Lamo-Espinosa, M. San-Julian, T. Jiménez-Solas, F. Lopez, S. Muntion, F. Sanchez-Guijo, A. Molero, J. Montoro, G. Serrano, A. Diaz-Mazkiaran, M. Lasaga, D. Gomez-Cabrero, M. Diez-Campelo, D. Valcarcel, M. Hernaez, J. P. Romero, F. Prosper, Uncovering perturbations in human hematopoiesis associated with healthy aging and myeloid malignancies at single-cell resolution. Elife 12 (2023).

51. C. López-Otín, M. A. Blasco, L. Partridge, M. Serrano, G. Kroemer, Hallmarks of aging: An expanding universe. Cell 186(2):243–278 (2023). 10.1016/j.cell.2022.11.001.

52. N. Guidi, G. Marka, V. Sakk, Y. Zheng, M. C. Florian, H. Geiger, An aged bone marrow niche restrains rejuvenated hematopoietic stem cells. Stem Cells 39, 1101–1106 (2021).

53. M. Kwon, B. S. Kim, S. Yoon, S. O. Oh, D. Lee, Hematopoietic Stem Cells and Their Niche in Bone Marrow. International Journal of Molecular Sciences 2024*, Vol.* 25, *Page* 6837 **25**, 6837 (2024).

54. C. Feng, H. Fan, R. Tie, S. Xin, M. Chen, Deciphering the evolving niche interactome of human hematopoietic stem cells from ontogeny to aging. Front Mol Biosci 11 (2024).

55. Y. H. Ho, R. del Toro, J. Rivera-Torres, J. Rak, C. Korn, A. García-García, D. Macías, C. González-Gómez, A. del Monte, M. Wittner, A. K. Waller, H. R. Foster, C. López-Otín, R. S. Johnson, C. Nerlov, C. Ghevaert, W. Vainchenker, F. Louache, V. Andrés, S. Méndez-Ferrer, Remodeling of Bone Marrow Hematopoietic Stem Cell Niches Promotes Myeloid Cell Expansion during Premature or Physiological Aging. Cell Stem Cell 25, 407–418.e6 (2019).

56. N. Yuan, W. Wei, L. Ji, J. Qian, Z. Jin, H. Liu, L. Xu, L. Li, C. Zhao, X. Gao, Y. He, M. Wang, L. Tang, Y. Fang, J. Wang, Young donor hematopoietic stem cells revitalize aged or damaged bone marrow niche by transdifferentiating into functional niche cells. Aging Cell 22 (2023).

57. Y. Han, S. Y. Kim, Endothelial senescence in vascular diseases: current understanding and future opportunities in senotherapeutics. Experimental & Molecular Medicine 2022 55:*1* 55, 1–12 (2023).

58. H. Wang, H. Xu, W. Chen, M. Cheng, L. Zou, Q. Yang, C. B. Chan, H. Zhu, C. Chen, J. Nie, B. Jiao, Rab13 Sustains Breast Cancer Stem Cells by Supporting Tumor-Stroma Cross-talk. Cancer Res 82, 2124–2140 (2022).

59. S. Winter, K. S. Götze, J. S. Hecker, K. H. Metzeler, B. Guezguez, K. Woods, H. Medyouf, A. Schäffer, M. Schmitz, R. Wehner, I. Glauche, I. Roeder, M. Rauner, L. C. Hofbauer, U. Platzbecker, Clonal hematopoiesis and its impact on the aging osteo-hematopoietic niche. Leukemia 2024 38:*5* 38, 936–946 (2024).

60. M. M. E. de Jong, Z. Kellermayer, N. Papazian, S. Tahri, D. Hofste op Bruinink, R. Hoogenboezem, M. A. Sanders, P. C. van de Woestijne, P. K. Bos, C. Khandanpour, J. Vermeulen, P. Moreau, M. van Duin, A. Broijl, P. Sonneveld, T. Cupedo, The multiple myeloma microenvironment is defined by an inflammatory stromal cell landscape. Nature Immunology 2021 22:*6* 22, 769–780 (2021).

61. Y. Gao, Y. Chi, Y. Chen, W. Wang, H. Li, W. Zheng, P. Zhu, J. An, Y. Duan, T. Sun, X. Liu, F. Xue, W. Liu, R. Fu, Z. Han, Y. Zhang, R. Yang, T. Cheng, J. Wei, L. Zhang, X. Zhang, Multi-omics analysis of human mesenchymal stem cells shows cell aging that alters immunomodulatory activity through the downregulation of PD-L1. Nat Commun 14 (2023).

62. S. Dudli, A. Karol, L. Giudici, I. Heggli, C. J. Laux, J. M. Spirig, F. Wanivenhaus, M. Betz, C. Germann, N. Farshad-Amacker, F. Brunner, O. Distler, M. Farshad, CD90-positive stromal cells associate with inflammatory and fibrotic changes in modic changes. Osteoarthr Cartil Open 4 (2022).

63. X. Yang, Y. Wang, V. Rovella, E. Candi, W. Jia, F. Bernassola, P. Bove, M. Piacentini, M. Scimeca, G. Sica, G. Tisone, A. Mauriello, L. Wei, G. Melino, Y. Shi, Aged mesenchymal stem cells and inflammation: from pathology to potential therapeutic strategies. Biol Direct 18, 1–12 (2023).

64. E. Maldonado, S. Morales-Pison, F. Urbina, A. Solari, Aging Hallmarks and the Role of Oxidative Stress. Antioxidants 2023, *Vol.* 12, *Page 651* **12**, 651 (2023).

65. M. Barilani, C. Lovejoy, R. Piras, A. Y. Abramov, L. Lazzari, P. R. Angelova, Age-related changes in the energy of human mesenchymal stem cells. J Cell Physiol 237, 1753–1767 (2022).

66. J. J. Mistry, K. A. Young, P. A. C. Díaz, I. F. Maestre, R. L. Levine, J. J. Trowbridge, Mesenchymal Stromal Cell Senescence Induced by Dnmt3a-Mutant Hematopoietic Cells is a Targetable Mechanism Driving Clonal Hematopoiesis and Initiation of Hematologic Malignancy. bioRxiv, 2024.03.28.587254 (2024).

67. How age affects human hematopoietic stem and progenitor cells and the strategies to mitigate aging. Exp Hematol 143, 104711 (2025).

68. C. Rastogi, H. T. Rube, J. F. Kribelbauer, J. Crocker, R. E. Loker, G. D. Martini, O. Laptenko, W. A. Freed-Pastor, C. Prives, D. L. Stern, R. S. Mann, H. J. Bussemaker, Counteracting age-related VEGF signaling insufficiency promotes healthy aging and extends life span. Science (1979) 373, E3692–E3701 (2021).

69. A. P. Kusumbe, S. K. Ramasamy, T. Itkin, M. A. Mäe, U. H. Langen, C. Betsholtz, T. Lapidot, R. H. Adams, Age-dependent modulation of vascular niches for haematopoietic stem cells. Nature 532, 380–384 (2016).

70. Z. Al-Yafeai, A. Yurdagul, J. M. Peretik, M. Alfaidi, P. A. Murphy, A. W. Orr, Endothelial FN (Fibronectin) deposition by α5β1 integrins drives atherogenic inflammation. Arterioscler Thromb Vasc Biol 38, 2601–2614 (2018).

71. L. Sudupe, E. Muiños-Lopez, A. R. Lopez-Perez, A. Vilas-Zornoza, S. Sarvide, P. Ripalda-Cemborain, P. Aguirre-Ruiz, P. S. Martin-Uriz, M. Larrayoz, L. Alvarez-Gigli, M. Abengozar-Muela, I. Cenzano, M. Cócera, J. Ruiz, I. S. González, A. Bantan, A. Kurowska, J. Ye, P. T. Newton, B. Paiva, J. R. Rodriguez-Madoz, V. Lagani, J. Tegner, B. Saez, J. A. Martinez-Climent, I. A. Calvo, D. Gomez-Cabrero, F. Prosper, Spatial Transcriptomics Reveals a Myeloma Cell Architecture with Dysfunctional T-Cell Distribution, Neutrophil Traps, and Inflammatory Signaling. bioRxiv, 2024.07.03.601833 (2024).

72. X. Xiao, C. Juan, T. Drennon, C. R. Uytingco, N. Vishlaghi, D. Sokolowskei, L. Xu, B. Levi, M. C. Sammarco, R. J. Tower, Spatial transcriptomic interrogation of the murine bone marrow signaling landscape. Bone Research 2023 11:*1* **11**, 1–13 (2023).

73. T. Stuart, A. Butler, P. Hoffman, C. Hafemeister, E. Papalexi, W. M. Mauck, Y. Hao, M. Stoeckius, P. Smibert, R. Satija, Comprehensive Integration of Single-Cell Data. Cell 177, 1888–1902.e21 (2019).

74. P. L. Germain, A. Lun, W. Macnair, M. D. Robinson, Doublet identification in single-cell sequencing data using scDblFinder. F1000Research 2021 10:*979* **10**, 979 (2021).

75. C. Hafemeister, R. Satija, Normalization and variance stabilization of single-cell RNA-seq data using regularized negative binomial regression. Genome Biol. 20, 1–15 (2019).

76. I. Korsunsky, N. Millard, J. Fan, K. Slowikowski, F. Zhang, K. Wei, Y. Baglaenko, M. Brenner, P. ru Loh, S. Raychaudhuri, Fast, sensitive and accurate integration of single-cell data with Harmony. Nat Methods 16, 1289–1296 (2019).

77. G. Finak, A. McDavid, M. Yajima, J. Deng, V. Gersuk, A. K. Shalek, C. K. Slichter, H. W. Miller, M. J. McElrath, M. Prlic, P. S. Linsley, R. Gottardo, MAST: A flexible statistical framework for assessing transcriptional changes and characterizing heterogeneity in single-cell RNA sequencing data. Genome Biol 16, 1–13 (2015).

78. G. Yu, L. G. Wang, Y. Han, Q. Y. He, ClusterProfiler: an R package for comparing biological themes among gene clusters. *OMICS A J*. Integr. Biol. 16, 284–287 (2012).

79. S. A. Miller, R. A. Policastro, S. Sriramkumar, T. Lai, T. D. Huntington, C. A. Ladaika, D. Kim, C. Hao, G. E. Zentner, H. M. O’Hagan, LSD1 and aberrant DNA methylation mediate persistence of enteroendocrine progenitors that support BRAF mutant colorectal cancer. Cancer Res 81, 3791 (2021).

80. D. Van De Vlekkert, E. Machado, A. D’azzo, Analysis of Generalized Fibrosis in Mouse Tissue Sections with Masson’s Trichrome Staining. doi: 10.21769/BioProtoc.3629 (2020).

