## Supplemental Material_figures for "Single-cell profiling reveals a novel RAB13+ endothelial subpopulation and profibrotic mesenchymal cells in the aged human bone marrow"

Itziar Cenzano *et al.*

**This PDF file includes:**

Figs. S1 to S10
Tables S1 to S12

**a**

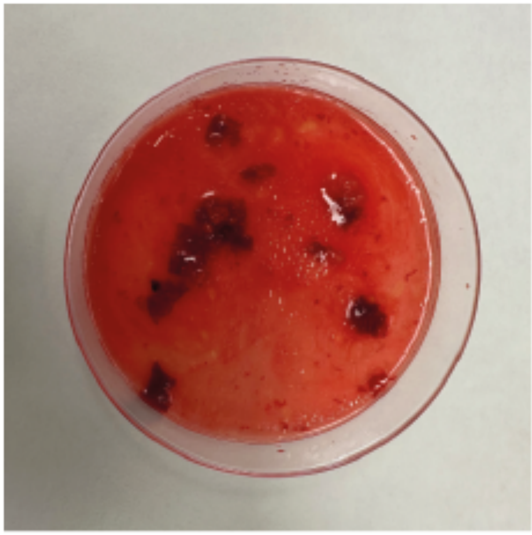

**b**

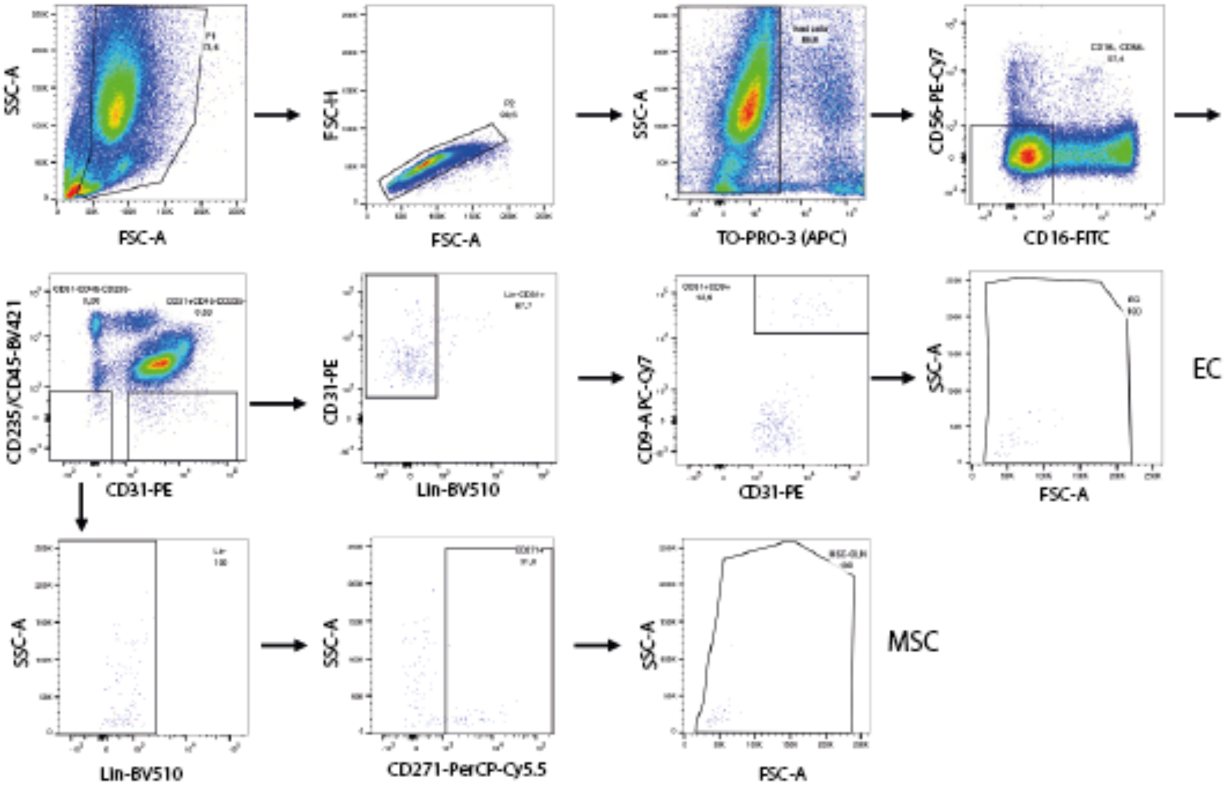
SSC-A
FSC-A
FSC-H
FSC-A
SSC-A
TO-PRO-3 (APC)
CD56-PE-Cy7
CD16-FITC
CD235/CD45-BV421
CD31-PE
CD31-PE
Lin-BV510
CD9-A PC-Cy7
CD31-PE
SSC-A
FSC-A
EC
SSC-A
Lin-BV510
SSC-A
CD271-PerCP-Cy5.5
SSC-A
FSC-A
MSC

**Fig. S1.**

**Cell sorting strategy for the isolation of human BM EC and MSC.** a. Representative image of processed human BM sample obtained from orthopedic hip replacement surgery. It contains a BM liquid fraction together with bone pieces. b. Sorting gating strategy for isolation of human BM EC (TO-PRO-3$^-$, CD16$^-$, CD56$^-$, CD45$^-$, CD235$^-$, Lin$^-$, CD31$^+$, CD9$^+$) and MSC (TO-PRO-3$^-$, CD16$^-$, CD56$^-$, CD45$^-$, CD235$^-$, CD31$^-$, Lin$^-$, CD271$^+$).

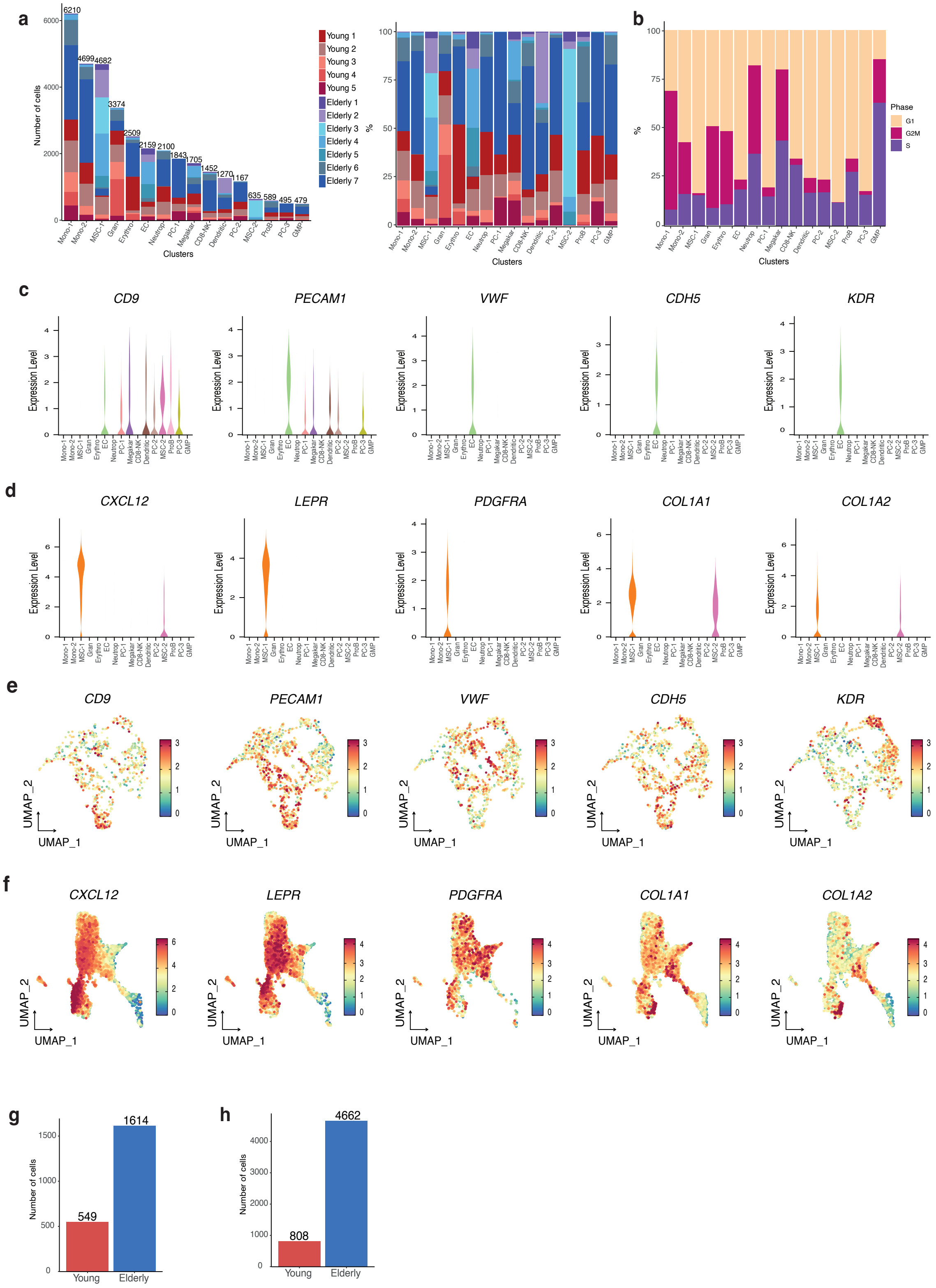
a
Number of cells
6210
4699
4682
3374
2509
2159
2100
1843
1705
1452
1270
1167
635
589
495
479
Clusters
Young 1
Young 2
Young 3
Young 4
Young 5
Elderly 1
Elderly 2
Elderly 3
Elderly 4
Elderly 5
Elderly 6
Elderly 7
%
Mono-1
Mono-2
MSC-1
Gran
Erythro
EC
Neutrop
PC-1
Megakar
CD8-NK
Dendritic
PC-2
MSC-2
ProB
PC-3
GMP
b
Phase
G1
G2M
S
c
CD9
PECAM1
VWF
CDH5
KDR
Expression Level
d
CXCL12
LEPR
PDGFRA
COL1A1
COL1A2
e
UMAP_2
UMAP_1
f
g
1614
549
Young
Elderly
h
4662
808

**Fig. S2**.

**scRNA-seq analysis of young and aged BM endothelial and stromal cells. a.** Left panel: bar plots showing the number of cells per cluster and dataset; Right panel: stacked bar plot representing the proportion of cells per cluster and dataset. **b**. Stacked bar plot representing the proportion of cells per cycle stage in each cluster. **c-d**. Violin plots showing the expression of canonical markers for EC (c) and MSC (d) per cluster. **e-f**. UMAP visualization of well-known markers for EC (e) and MSC (f). **g-h**. Bar plot depicting the total number of EC (g) and MSC (h) per age group (young and elderly).

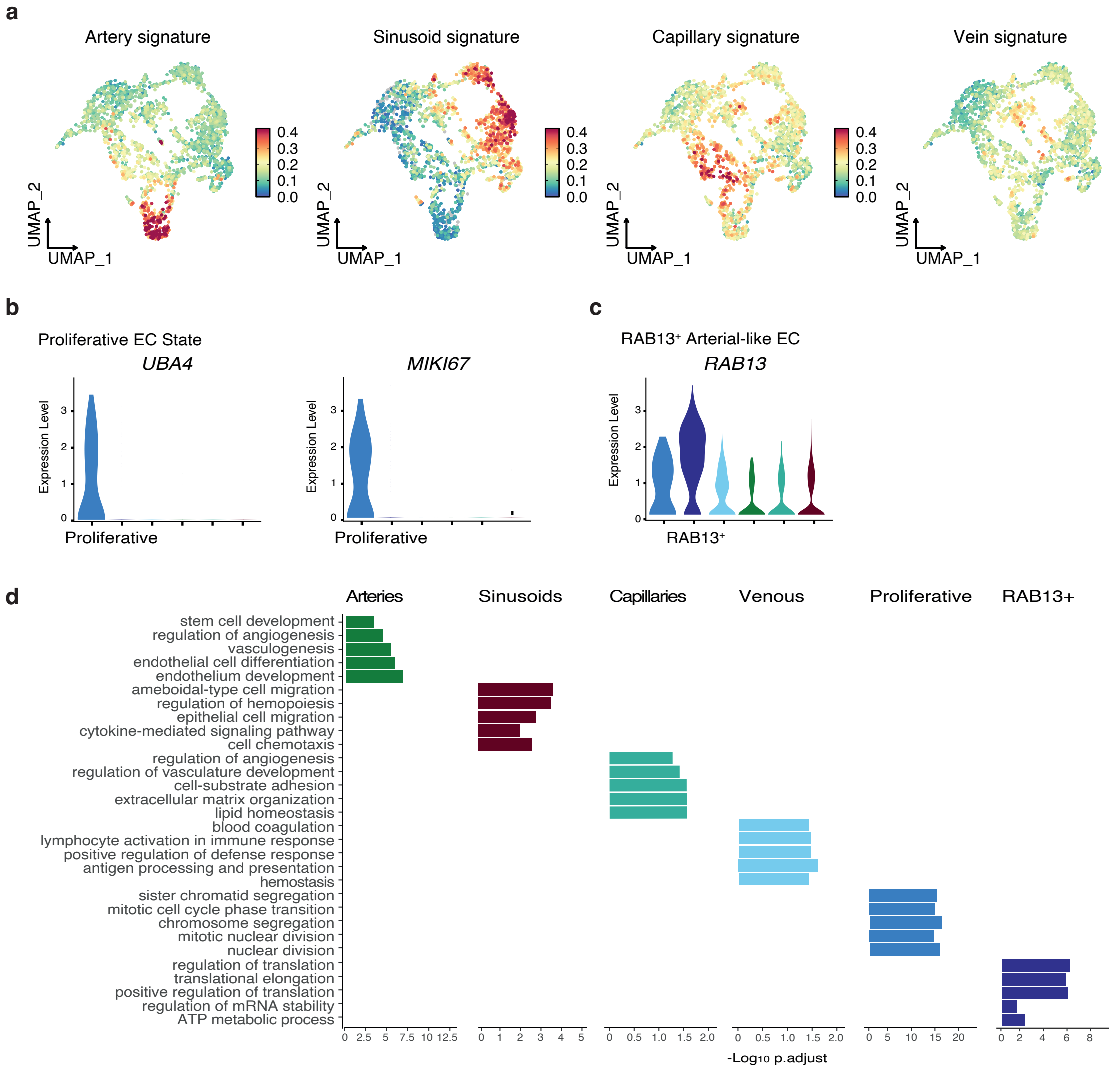


**Fig. S3.**

**Signatures and markers defining the vascular beds in the human BM endothelial compartment. a**. UMAP visualization of artery, sinusoid, capillary, and vein signature scores. **b**. Violin plot displaying the expression of cell cycle-related genes. **c**. Violin plot showing the expression distribution of *RAB13*$^{+}$-arterial-like EC in all EC. **d**. Significant gene sets derived from the GO ORA conducted with the markers defining each vascular state.

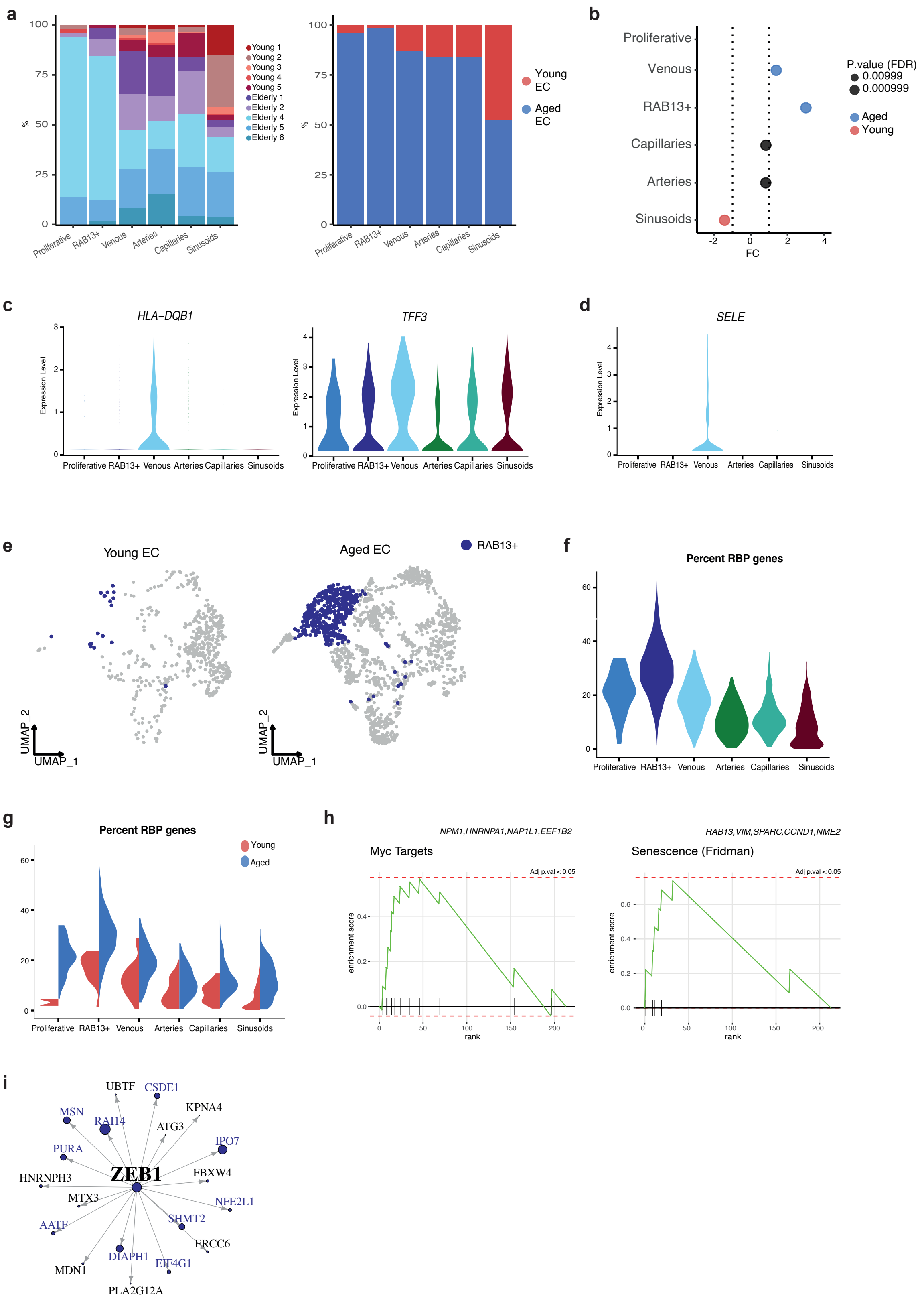
a
Young 1
Young 2
Young 3
Young 4
Young 5
Elderly 1
Elderly 2
Elderly 4
Elderly 5
Elderly 6
Young EC
Aged EC
Proliferative
RAB13+
Venous
Arteries
Capillaries
Sinusoids
b
P.value (FDR)
0.00999
0.000999
Aged
Young
FC
c
HLA−DQB1
TFF3
Expression Level
d
SELE
e
Young EC
Aged EC
RAB13+
UMAP_1
UMAP_2
f
Percent RBP genes
g
Percent RBP genes
Young
Aged
h
NPM1,HNRNPA1,NAP1L1,EEF1B2
Myc Targets
RAB13,VIM,SPARC,CCND1,NME2
Senescence (Fridman)
Adj p.val < 0.05
enrichment score
rank
i
ZEB1
UBTF
CSDE1
KPNA4
MSN
RAI14
ATG3
IPO7
PURA
HNRNPH3
FBXW4
MTX3
NFE2L1
AATF
SHMT2
ERCC6
DIAPH1
EIF4G1
MDN1
PLA2G12A

**Fig. S4.**

**Characterization of the aged endothelial compartment. a**. Stacked bar plots representing the proportion of cells per individual (left panel) and age group (right panel) in each cluster. **b**. Relative differences in cell proportions for each vascular state, comparing young and aged EC. Red and blue represent clusters statistically significant (FDR < 0.05 and absolute log2 fold change > 1) in young and aged EC, respectively. Larger log2 fold changes indicate a higher proportion of cells. **c.** Violin plot showing the expression of genes upregulated in the venous cluster. **d.** Violin plot displaying the expression of *SELE* among the vascular states. **e**. UMAP visualization of the distribution of $RAB13^+$ cells in young (left) and aged (right) EC. **f**. Violin plot displaying the percentage of ribosomal binding proteins (RBP) per EC cluster. **g**. Split violin plots showing the percentage of RBP in EC clusters split by age group. **h**. GSEA plot showing the enrichment of "Myc targets" and "Senescence" terms in $RAB13^+$ EC. **i**. Network of ZEB1 regulon enriched in $RAB13^+$ EC.

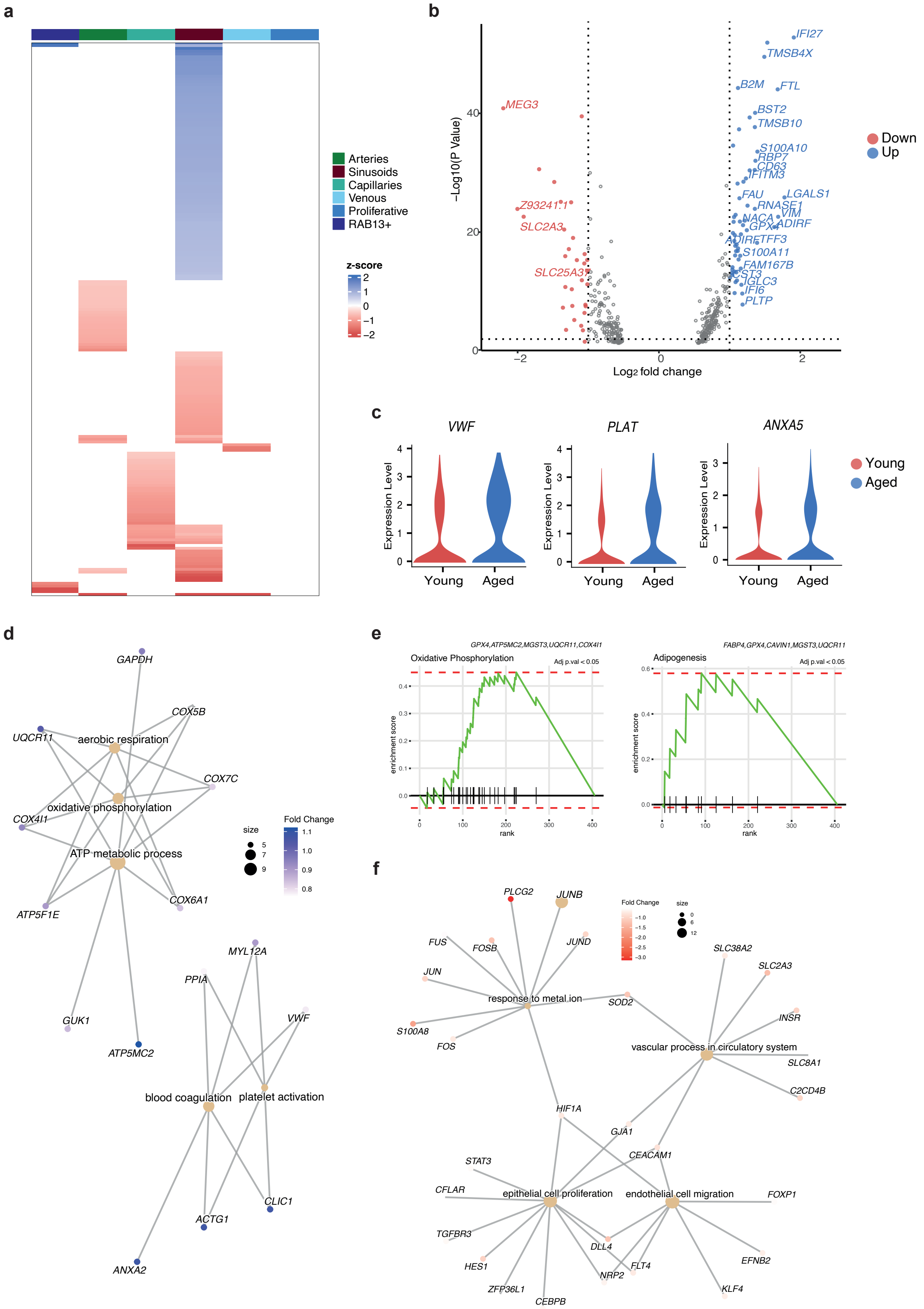
a
Arteries
Sinusoids
Capillaries
Venous
Proliferative
RAB13+
z-score
2
1
0
−1
−2
b
IFI27
TMSB4X
B2M
FTL
BST2
TMSB10
S100A10
RBP7
CD63
IFITM3
FAU
LGALS1
RNASE1
VIM
NACA
GPX4
ADIRF
TFF3
S100A11
FAM167B
CST3
IGLC3
IFI6
PLTP
MEG3
Z93241.1
SLC2A3
SLC25A37
−Log10(P Value)
Log2 fold change
Down
Up
c
VWF
PLAT
ANXA5
Expression Level
Young
Aged
d
GAPDH
COX5B
UQCR11
aerobic respiration
COX7C
oxidative phosphorylation
COX4I1
ATP metabolic process
COX6A1
ATP5F1E
GUK1
ATP5MC2
MYL12A
PPIA
VWF
blood coagulation
platelet activation
CLIC1
ACTG1
ANXA2
size
Fold Change
e
GPX4,ATP5MC2,MGST3,UQCR11,COX4I1
Oxidative Phosphorylation
Adj p.val < 0.05
FABP4,GPX4,CAVIN1,MGST3,UQCR11
Adipogenesis
enrichment score
rank
f
PLCG2
JUNB
FUS
FOSB
JUND
JUN
response to metal ion
SOD2
S100A8
FOS
SLC38A2
SLC2A3
INSR
vascular process in circulatory system
SLC8A1
C2CD4B
HIF1A
GJA1
CEACAM1
STAT3
CFLAR
epithelial cell proliferation
endothelial cell migration
FOXP1
TGFBR3
DLL4
HES1
FLT4
NRP2
EFNB2
ZFP36L1
CEBPB
KLF4

**Fig. S5.**

**Transcriptional remodeling of sinusoids in the aged BM EC. a.** Summary heatmap of the number and effect size of all age-DEGs (FDR<0.05; abs(log2FC)>0.5) identified within each vascular state. Color represents the z-score of the log2FC. **b**. Volcano plot of the DEGs between young and aged sinusoids. The y-axis represents the -log10(p-value), and the x-axis represents the log2(fold change). The color of the dot denotes the age group for which DEGs were detected, with grey dots representing non-significant genes. **c**. Violin plots showing the expression of aging-related prothrombotic and matrix-associated genes upregulated in aged sinusoids. **d**. Cnetplot showing the links between genes and biological processes upregulated in aged sinusoids. Node size reflects the number of significantly enriched genes in the node and colors the $\log_2$ Fold change expression of each gene. **e**. GSEA plot of "Oxidative phosphorylation" and "Adipogenesis" terms significantly enriched in aged sinusoids. **f**. Cnetplot showing the relationship among individual GO terms and genes downregulated in aged sinusoids. Node size indicates the number of significantly enriched genes in the node and colors the $\log_2$ Fold change expression of each gene.

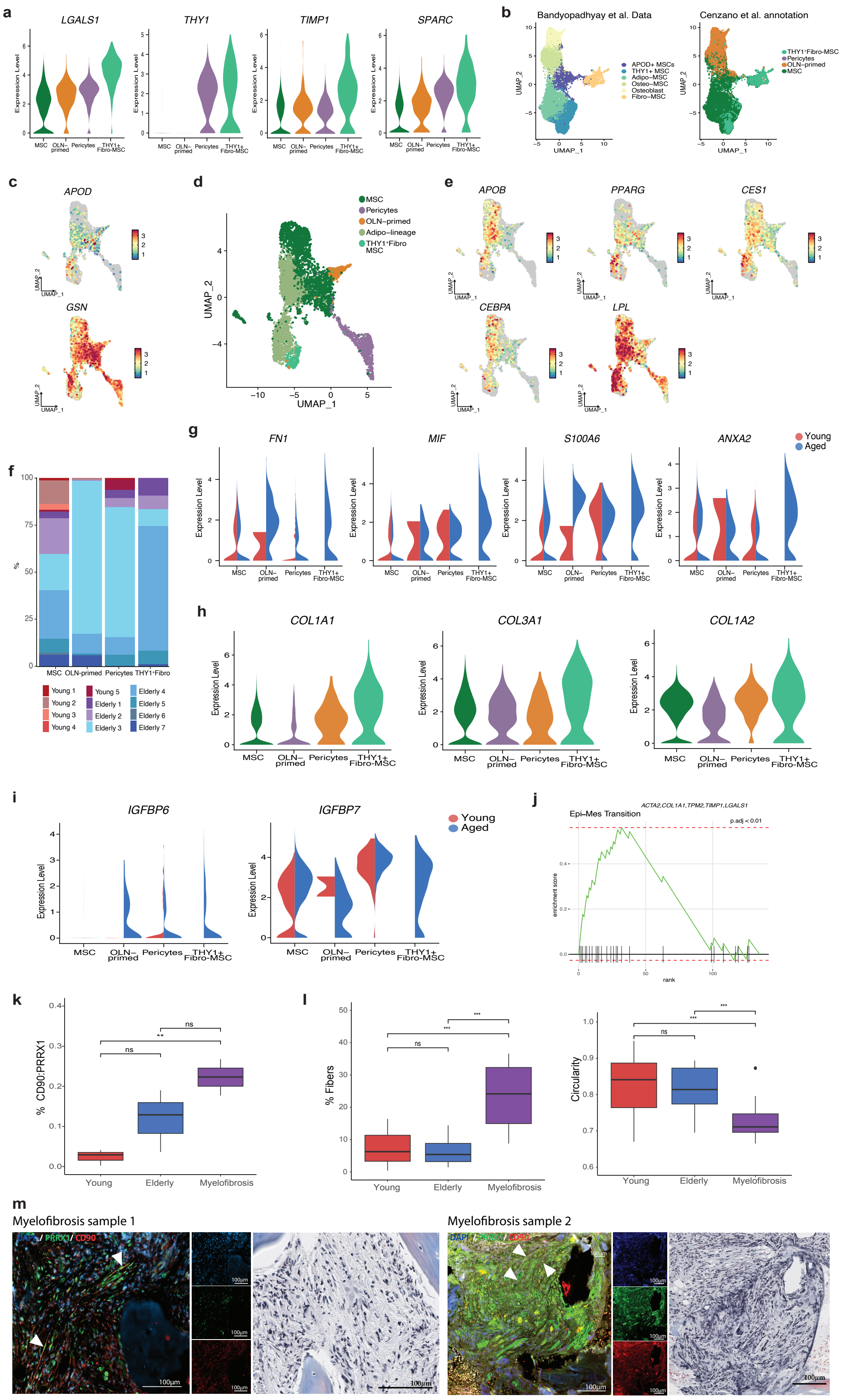
a
LGALS1
THY1
TIMP1
SPARC
Expression Level
MSC
OLN–primed
Pericytes
THY1+ Fibro-MSC
b
Bandyopadhyay et al. Data
Cenzano et al. annotation
APOD+ MSCs
THY1+ MSC
Adipo–MSC
Osteo–MSC
Osteoblast
Fibro–MSC
THY1+Fibro-MSC
Pericytes
OLN–primed
MSC
UMAP_1
UMAP_2
c
APOD
GSN
d
MSC
Pericytes
OLN–primed
Adipo–lineage
THY1+Fibro MSC
e
APOB
PPARG
CES1
CEBPA
LPL
f
%
MSC
OLN-primed
Pericytes
THY1+Fibro
Young 1
Young 2
Young 3
Young 4
Young 5
Elderly 1
Elderly 2
Elderly 3
Elderly 4
Elderly 5
Elderly 6
Elderly 7
g
FN1
MIF
S100A6
ANXA2
Young
Aged
h
COL1A1
COL3A1
COL1A2
i
IGFBP6
IGFBP7
Young
Aged
j
ACTA2,COL1A1,TPM2,TIMP1,LGALS1
Epi–Mes Transition
p.adj < 0.01
enrichment score
rank
k
% CD90:PRRX1
Young
Elderly
Myelofibrosis
ns
**
l
% Fibers
Circularity
***
m
Myelofibrosis sample 1
Myelofibrosis sample 2
DAPI / PRRX1 / CD90
100µm

**Fig. S6.**

**Further transcriptional profiling of THY1+Fibro-MSC. a**. Violin plots showing the marker expression of THY1$^+$Fibro-MSC cluster. **b**. Left: UMAP visualization of the Bandyopadhyay et al.[1] dataset. Colors denote the identified stromal subtypes. Right: UMAP projection showing the distribution of stromal cell type annotations from our study in the Bandyopadhyay et al.[1] dataset. **c**. UMAP visualization of APOD and GSN genes defining APOD$^+$GSN$^{high}$ MSC described in Bandyopadhyay et al.[1] dataset. **d**. UMAP plot illustrating the distribution of stromal cell subtypes, including adipo-lineage clusters. **e**. UMAP visualization of the expression of adipogenic-related genes upregulated in adipo-lineage cells. **f**. Stacked bar plots representing the proportion of cells per individual in each stromal subpopulation. **g**. Split violin plots showing the expression of aging-related regulators upregulated in the THY1$^+$Fibro-MSC cluster split by age. **h**. Violin plot showing the expression of collagen-associated genes related to TGF-β signaling pathway upregulated in the THY1$^+$Fibro-MSC cluster. **i**. Aged-split violin plots showing the expression of *IGFBP6* and *IGFBP7* in MSC clusters. **j**. GSEA plot of "Epithelial-Mesenchymal (Epi-Mes) Transition" term significantly enriched in THY1$^+$Fibro-MSC cluster. **k.** Quantification of THY1$^+$ cells (CD90$^+$) (red), MSC (PRRX1$^+$) (green) and THY1$^+$MSC (coexpression CD90$^+$ and PRRX1$^+$) in young, elderly, and myelofibrosis (MF) samples. Bars represent the mean ± SEM. ns: not significant. **l.** Quantification of fibrotic tissue area in young, elderly, and MF samples. The fibrotic tissue area was measured in Masson's trichrome-stained images by two approximations. The percent of fibrotic tissue area/whole tissue area and the circularity of the cells. Bars represent the mean ± SEM. ns: not significant. **m.** Left panel: IF staining of THY1$^+$ Fibro stromal cells (CD90$^+$) (red), (PRRX1$^+$) (green), and nucleus (DAPI) (blue) in FFPE biopsy samples from MF patients. Scale bars: 100 µm. Right panel: Masson-Trichrome staining for fibrotic tissue of FFPE biopsies from the same tissue area. White arrows indicate THY1$^+$ Fibro MSC (PRRX1+CD90+).

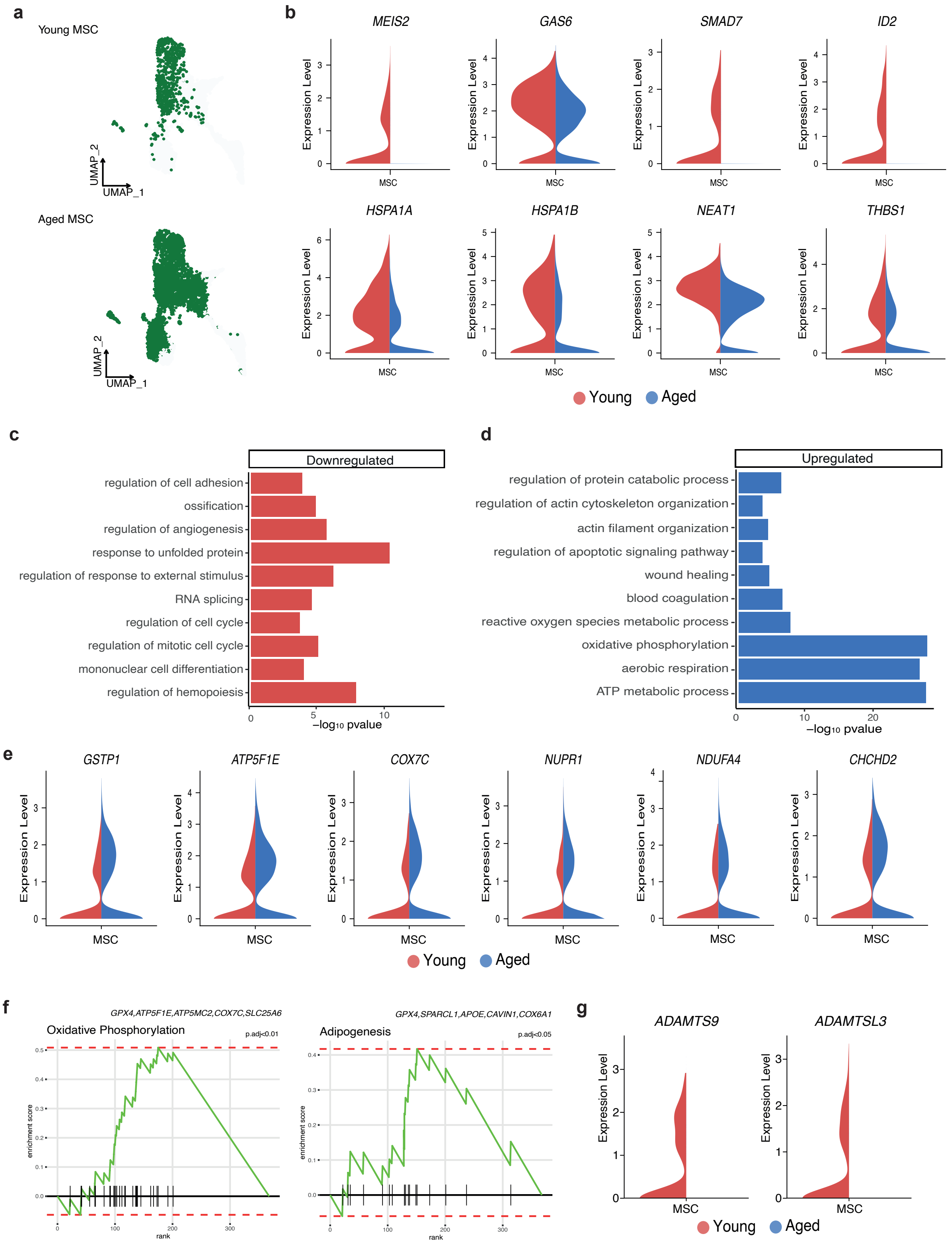
a
Young MSC
Aged MSC
UMAP_1
UMAP_2
b
MEIS2
GAS6
SMAD7
ID2
HSPA1A
HSPA1B
NEAT1
THBS1
Expression Level
MSC
Young
Aged
c
Downregulated
regulation of cell adhesion
ossification
regulation of angiogenesis
response to unfolded protein
regulation of response to external stimulus
RNA splicing
regulation of cell cycle
regulation of mitotic cell cycle
mononuclear cell differentiation
regulation of hemopoiesis
−log10 pvalue
d
Upregulated
regulation of protein catabolic process
regulation of actin cytoskeleton organization
actin filament organization
regulation of apoptotic signaling pathway
wound healing
blood coagulation
reactive oxygen species metabolic process
oxidative phosphorylation
aerobic respiration
ATP metabolic process
−log10 pvalue
e
GSTP1
ATP5F1E
COX7C
NUPR1
NDUFA4
CHCHD2
Expression Level
MSC
Young
Aged
f
GPX4,ATP5F1E,ATP5MC2,COX7C,SLC25A6
Oxidative Phosphorylation
p.adj<0.01
GPX4,SPARCL1,APOE,CAVIN1,COX6A1
Adipogenesis
p.adj<0.05
enrichment score
rank
g
ADAMTS9
ADAMTSL3
Expression Level
MSC
Young
Aged

**Fig. S7.**

**Age-dependent transcriptional changes in MSC. a**. UMAP visualization of MSC in young (top) and aged (bottom). **b**. Split violin plots showing the expression of down-regulated genes in aged MSC split by age group. **c-d**. Bar charts of enriched GO terms from ORA (p-value < 0.05) comparing DEGs within young and aged MSC. The horizontal axis represents the -log10 of p-values. (c) represents downregulated terms and (d) upregulated terms in aged MSC. **e**. Split violin plots showing the expression of oxidative metabolism-related genes upregulated in aged MSC. **f**. GSEA plot of "Oxidative phosphorylation" and "Adipogenesis" terms significantly enriched in aged MSC. **g**. Split violin plots showing the downregulation of *ADAMTS9* and *ADAMTSL3* in aged MSC.

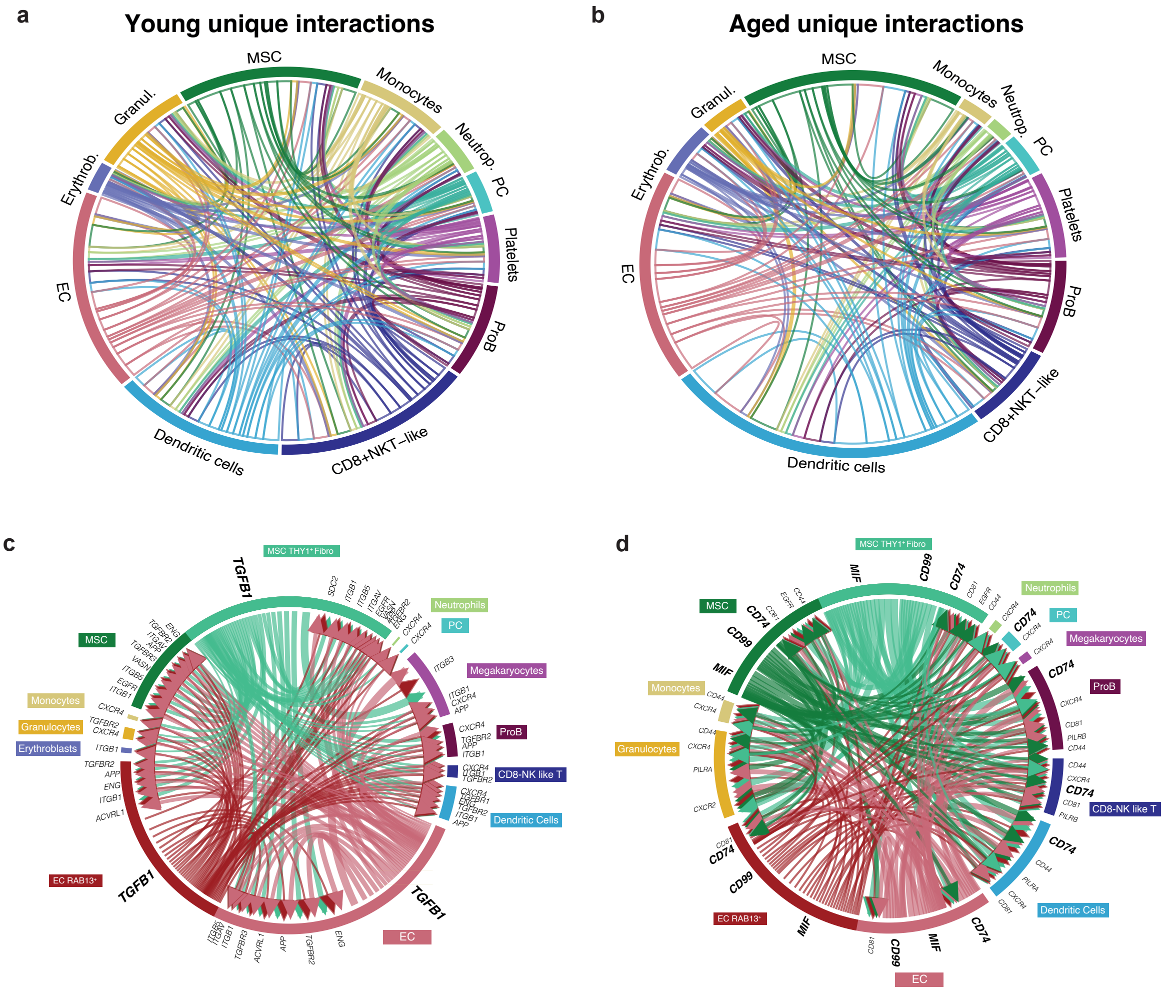
a
Young unique interactions
MSC
Monocytes
Neutrop.
PC
Platelets
ProB
CD8+NKT−like
Dendritic cells
EC
Erythrob.
Granul.
b
Aged unique interactions
MSC
Monocytes
Neutrop.
PC
Platelets
ProB
CD8+NKT−like
Dendritic cells
EC
Erythrob.
Granul.
c
MSC THY1+ Fibro
TGFB1
Neutrophils
PC
Megakaryocytes
ProB
CD8-NK like T
Dendritic Cells
EC
EC RAB13+
MSC
Monocytes
Granulocytes
Erythroblasts
d
MSC THY1+ Fibro
MIF
CD99
CD74
Neutrophils
PC
Megakaryocytes
ProB
CD8-NK like T
Dendritic Cells
EC
EC RAB13+
MSC
Monocytes
Granulocytes

**Fig. S8.**

**Additional information about the remodeling of the BM interactome during aging. a-b**. Chord diagrams displaying unique interactions specific to young (a) and aged (b) BM microenvironment cells. **c**. Chord diagrams showing interactions involving *TGFB1* signaling in the aged BM. Colors and widths represent the signal senders and the strength of interactions, respectively. **d**. Chord diagrams showing interactions through *MIF*, *CD74*, and *CD99* ligands in the aged BM. Colors and widths represent the signal senders and the strength of interactions, respectively.

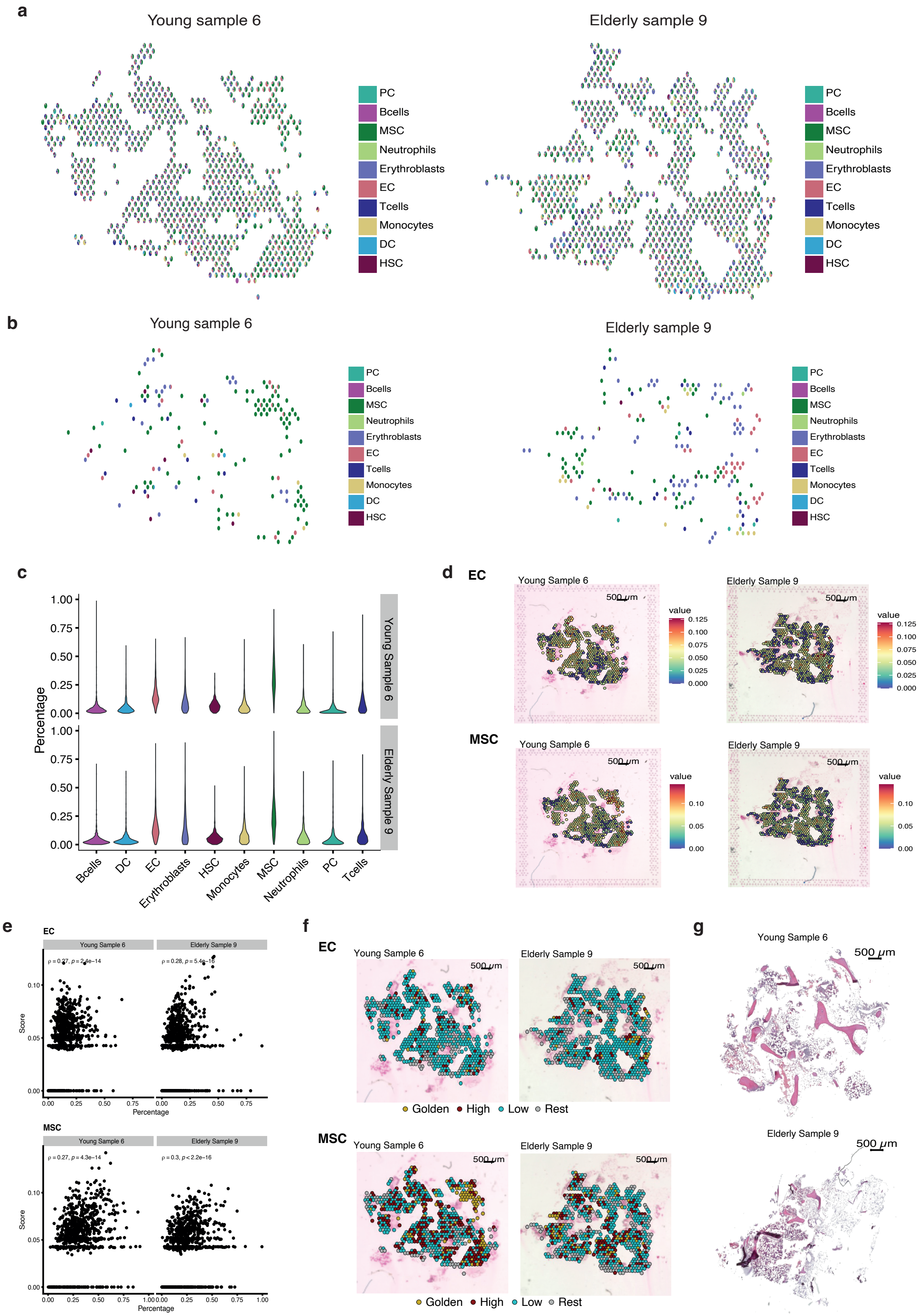
a
Young sample 6
Elderly sample 9
PC
Bcells
MSC
Neutrophils
Erythroblasts
EC
Tcells
Monocytes
DC
HSC
b
Young sample 6
Elderly sample 9
c
Percentage
Young Sample 6
Elderly Sample 9
Bcells
DC
EC
Erythroblasts
HSC
Monocytes
MSC
Neutrophils
PC
Tcells
d
EC
MSC
Young Sample 6
Elderly Sample 9
500 μm
value
e
Score
Percentage
f
Golden
High
Low
Rest
g
Young Sample 6
Elderly Sample 9
500 μm

**Fig. S9.**

**Additional information about spatial transcriptomics analysis of the human BM. a**. Pie charts illustrating the proportion of each cell type contributing to the transcriptomic signature of each spot in young (left) and elderly (right) BM samples from deconvolution analysis. **b**. Pie charts illustrate the proportion of each cell type that contributes significantly to the transcriptomic signature of each spot in young (left) and elderly (right) BM samples. **c**. Cell type proportions per spot derived from deconvolution analysis using the Bandyopadhyay et al.[1] dataset as a reference. **d**. Spatial distribution pattern of the signature score for EC (top panels) and MSC (bottom panels) in young (left side) and elderly (right side) BM samples. **e**. Correlation between cell-type proportions obtained through deconvolution (percentage) and the signature scores of EC (top) and MSC (bottom) in young (left side) and elderly (right side) BM samples. **f**. Spatial distribution of "golden", "high", "low", and "rest" spots based on the top-ranking overlap between deconvolution and spot signature analyses (detailed in the methods section) in young (left side) and elderly (right side) BM samples using EC (top panels) and MSC (bottom panels) as examples. **g**. H&E staining of young (upper panel) and elderly (bottom panel) BM samples.

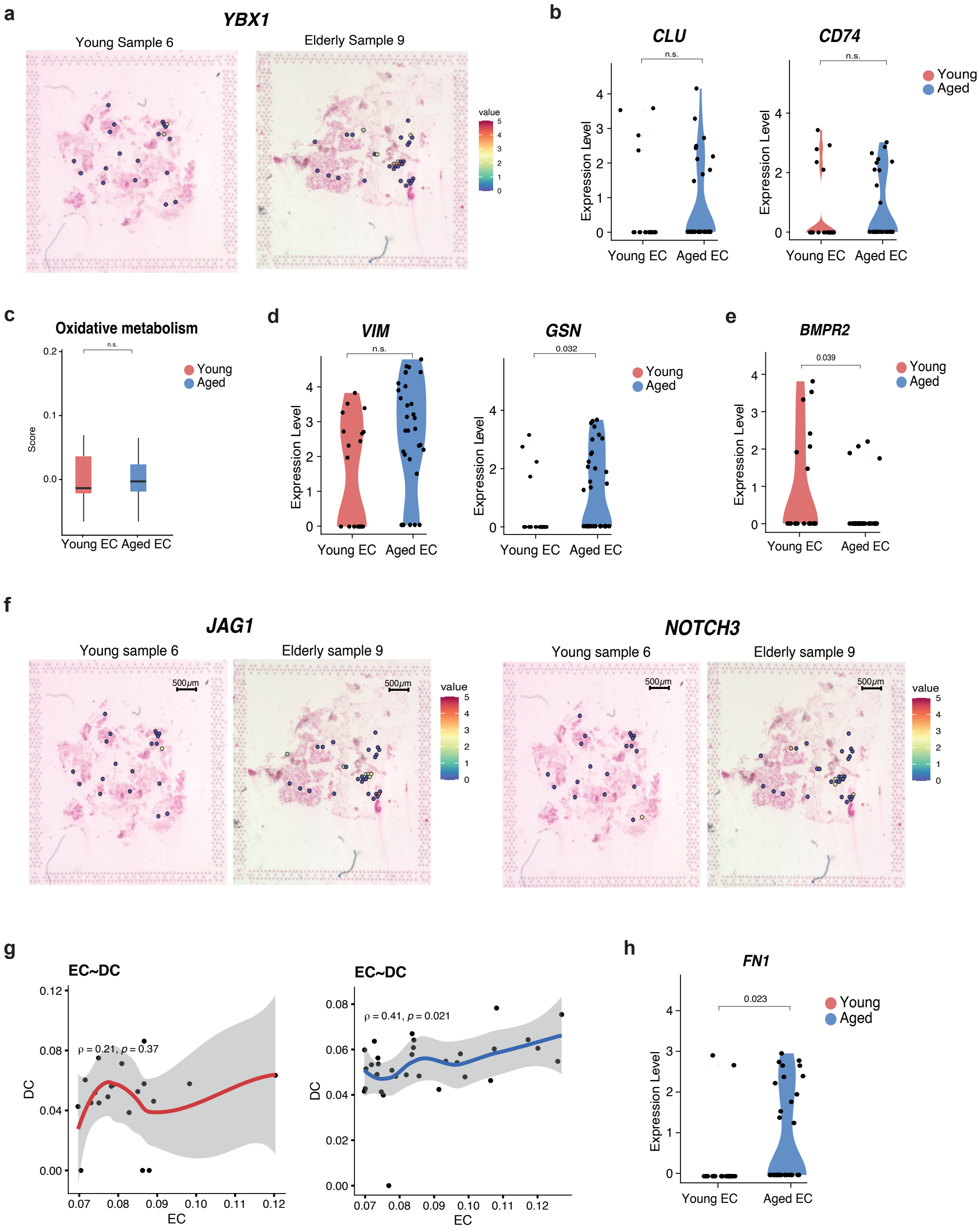
a
YBX1
Young Sample 6
Elderly Sample 9
value
b
CLU
CD74
n.s.
Young
Aged
Expression Level
Young EC
Aged EC
c
Oxidative metabolism
Score
d
VIM
GSN
0.032
e
BMPR2
0.039
f
JAG1
NOTCH3
Young sample 6
Elderly sample 9
500μm
g
EC~DC
ρ = 0.21, p = 0.37
ρ = 0.41, p = 0.021
DC
EC
h
FN1
0.023

**Fig. S10.**

**Additional information about the spatial expression patterns of aging-related gene expression changes in EC. a**. Spatial expression patterns of the transcription factor *YBX1* in young (left side) and elderly (right side) BM samples. **b**. Violin plots illustrating the spatial expression levels of CLU and CD74 genes in young (red) and aged (blue) EC. **c**. Box plots showing the scores for oxidative metabolism genes set in young (red) and aged (blue) EC. **d**. Violin plots illustrating the spatial expression levels of VIM and GSN genes in young (red) and aged (blue) EC **e**. Violin plots showing *BMPR2* spatial gene expression levels in young (red) and aged (blue) EC. **f**. Spatial expression patterns of the *JAG1-NOTCH3* L-R pair in EC golden spots of young (left panels) and elderly (right panels) BM samples. **g**. Plots showing the correlation between EC signature and DC signature of EC-labelled spots in young (left-red) and elderly (right-blue) samples. **h**. Violin plots of *FN1* spatial expression levels in young (red) and aged (blue) EC.
